# Dual apical methyltransferases orchestrate motility initiation in apicomplexan parasites

**DOI:** 10.1101/2025.11.05.686696

**Authors:** Peipei Qin, Thrishla Kumar, Oliwia Koczy, Wei Li, Ignasi Forne, Simone Mattei, Elena Jimenez-Ruiz

## Abstract

Apicomplexan parasites such as *Toxoplasma gondii* initiate motility through rapid, spatially confined cytoskeletal activation at their apical end. While calcium-, lipid-, and kinase-based signalling pathways have been partially elucidated, how these cues are translated into mechanical force remains unclear. Here, we uncover a dual methyltransferase mechanism that orchestrates this process. We identify PCKMT, a preconoidal ring–associated lysine methyltransferase, as an essential upstream regulator of motility. PCKMT anchors the actin nucleator Formin-1 (FRM1) at the conoid, enabling conoid protrusion and F-actin assembly. Loss of PCKMT abolishes FRM1 recruitment, blocks conoid extrusion, and arrests invasion and egress despite preserved conoid structure. In contrast, the apical methyltransferase AKMT, previously linked to motility through recruitment of the glideosome-associated connector (GAC), acts downstream, disengaging from the conoid upon activation and likely promoting GAC-dependent force transmission. PCKMT depletion prevents AKMT translocation, revealing that actin assembly and lysine methylation are mechanistically coupled.

Together, these findings define a two-step methylation checkpoint that coordinates actin nucleation with force propagation, uncovering methylation as a central regulatory axis for motility initiation in apicomplexan parasites.

## Main

Apicomplexan parasites such as *Toxoplasma gondii* rely on tightly coordinated cytoskeletal changes to invade and exit host cells. At the apex, the conoid complex and its preconoidal rings (PCRs) have emerged as regulatory hubs for motility initiation. Once enigmatic, these rings are now recognised as platforms that concentrate effectors essential for invasion and egress^1^. For example, the Conoid Gliding Protein (CGP) localises to the PCRs and is required for their integrity; parasites lacking CGP lose the rings and become non-motile^2, 3^. Thus, the PCRs act as architectural and regulatory hubs where motility factors assemble to ensure that gliding begins at the right time and place^4^.

Motility initiation is also governed by intracellular signalling. In *T. gondii*, the cGMP–PKG cascade elevates Ca²⁺ and triggers microneme secretion^5,6^. This is followed by conoid extrusion and F-actin nucleation at the apex by Formin-1 (FRM1), an actin regulator essential for invasion^7,8^. Newly formed actin filaments (F-actin) are then linked to surface adhesins by the glideosome-associated connector (GAC), thereby transmitting force for directed motion^9,10^. Despite this well-defined kinase and actin network, the contribution of post-translational modifications, particularly lysine methylation, to motility initiation remains largely unexplored. Although direct functional studies are limited, there is strong precedent that methylation modulates cytoskeletal dynamics and motility in other systems. For example, SETD2-mediated methylation of actin affects cell migration^11^. In addition, independent studies have identified methylated proteins in motile cilia and flagella^12,13^, suggesting a broader link between methylation and cytoskeletal dynamics.

In apicomplexan parasites, the apical lysine methyltransferase AKMT represents one of the few methylation enzymes directly implicated in motility control. It localises to the conoid and is essential for parasite egress and invasion; upon motility induction, AKMT rapidly translocates from the conoid to the cytosol^14,15^. This dynamic behaviour suggests a regulatory role at the apex, where methylation may act as a reversible switch coupling conoid extrusion to actin assembly. Here, we identify a second SET-domain methyltransferase, PCKMT (PreConoidal Ring-associated Lysine MethylTransferase), that localises to the preconoidal rings (PCRs) and acts upstream of AKMT to trigger motility initiation. Using conditional knockouts, high-resolution imaging, and proteomics, we demonstrate that PCKMT recruits the actin nucleator Formin-1 (FRM1) to the conoid to enable conoid protrusion and actin filament assembly. Loss of PCKMT abolishes FRM1 recruitment, blocks conoid protrusion, and prevents the actin-dependent disengagement of AKMT from the apex. Conversely, a catalytically inactive PCKMT mutant supports partial FRM1 recruitment but fails to restore motility, demonstrating that methyltransferase activity is essential. Together, these findings reveal a dual-methyltransferase checkpoint at the conoid: PCKMT licenses actin assembly by anchoring FRM1, thereby enabling AKMT-driven remodelling and GAC recruitment. This two-step methylation logic synchronises actin nucleation with force transmission to initiate motility.

## Results

### PCKMT is a conserved apical methyltransferase associated with the preconoidal rings

In previous studies, we identified several apical complex components essential for parasite egress^2^. Among them, CGP was shown to be critical for maintaining the integrity of the PCRs in mature *T. gondii* tachyzoites^3^. CGP depletion led to loss of the PCRs and impaired retention of key motility factors at mature conoid complexes, including the actin nucleator FRM1 (Fig. 1a). In the same study, a proximity-labelling assay identified a previously uncharacterised SET-domain methyltransferase as a potential CGP interactor. This protein is localised exclusively to the PCRs, and its stable association depended on CGP. Immunofluorescence and high-resolution imaging confirmed that this methyltransferase, hereafter referred to as PCKMT (PreConoidal Ring-associated Lysine MethylTransferase), resides at the preconoidal rings, positioned immediately above the conoid body motor myosin MyoH^16^ (Fig. 1b). Structural prediction using AlphaFold3 revealed that PCKMT (∼175 kDa) adopts a multidomain architecture composed of conserved TPR, ankyrin, and SET domains, followed by a tandem proline-rich region at the C-terminus (Supplemenatary Fig. 1a,b).

**Fig. 1:**
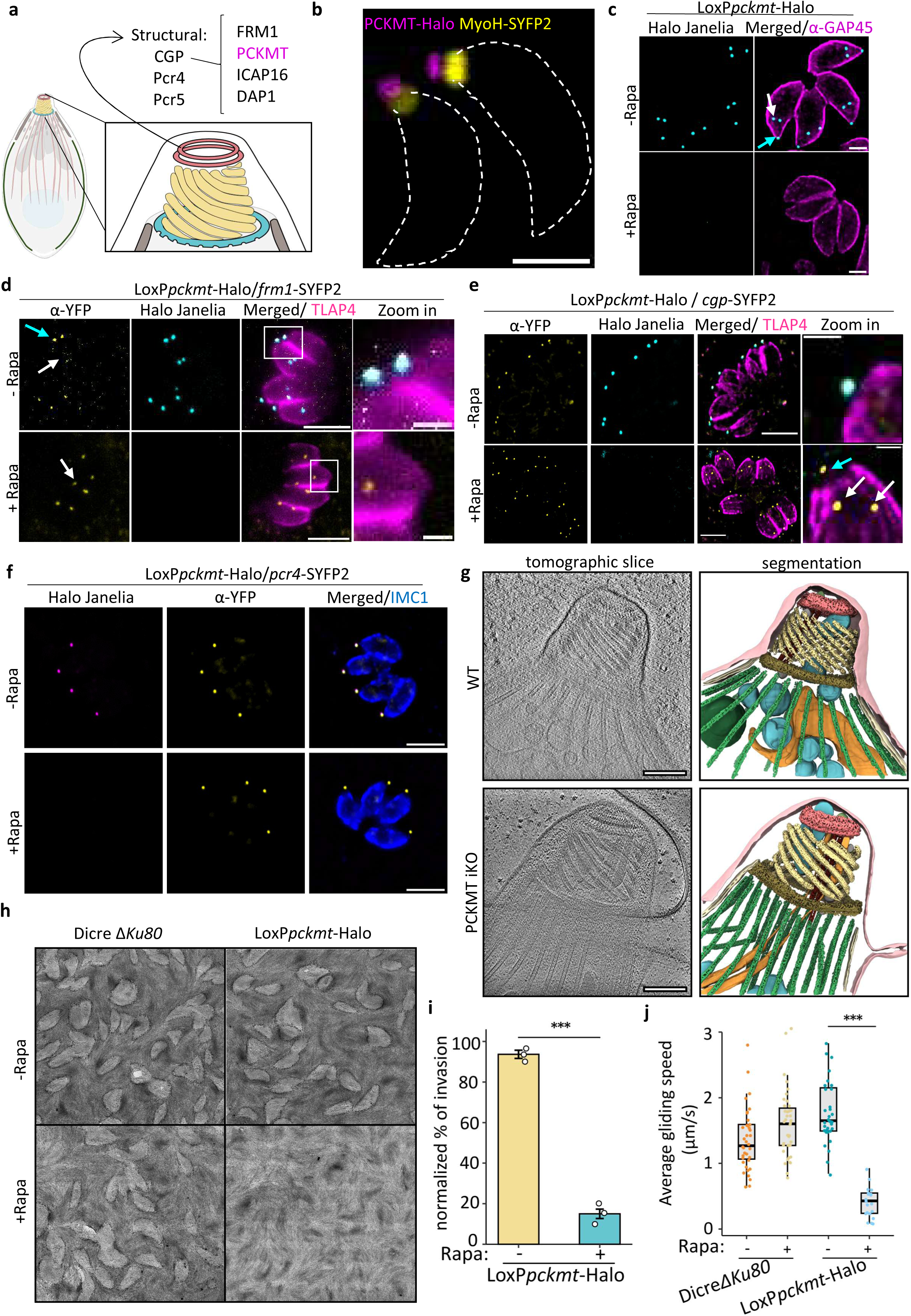
PCKMT localises to the preconoidal rings, anchors FRM1, and is essential for motility in *T. gondii*. **(a)** Schematic overview of the preconoidal ring (PCR) network. Structural components (CGP, Pcr4, and Pcr5) form the PCR scaffold, while CGP supports the recruitment of a subset of regulatory proteins (FRM1, ICAP16, DAP1, and PCKMT – shown in magenta-) at the preconoidal rings. **(b)** STED super-resolution image of a *T. gondii* tachyzoite showing that PCKMT (magenta) forms a discrete ring positioned apical to MyoH (yellow), consistent with preconoidal ring localisation. Scale bar, 1 µm. **(c)** Localisation of PCKMT at the conoid of mature parasites (cyan arrow) and developing daughter cells (white arrow) in non-induced parasites. Upon rapamycin induction, PCKMT signal is lost. Scale bar, 2 µm; representative of n = 3. **(d–f)** Endogenous tagging of FRM1 (d), CGP (e), and Pcr4 (f) in the conditional loxP*pckmt*-Halo strain shows that FRM1 signal is markedly reduced at the conoid upon PCKMT depletion, whereas CGP and Pcr4 remain unaffected, indicating a unidirectional dependency. Cyan arrow: mature conoids; white arrows: daughter cell conoids. **(g)** Tomographic slices and corresponding segmentations from cryo-electron tomograms of wild-type (WT) and PCKMT-depleted parasites (PCKMT iKO). The preconoidal rings of PCKMT-iKO parasites appear intact and morphologically unchanged. Segmented cellular structures are colour-coded: bright pink, plasma membrane; dark pink, preconoidal rings (PCRs); yellow, conoid; khaki, apical polar ring (APR); green, microtubules (MTs); orange, rhoptries; turquoise, micronemes; beige, inner membrane complex (IMC); dark red, intraconoidal microtubules (ICMTs); grey, microtubule-associated vesicles; light green, apical vesicle (AV); dark green, dense granule (observed only in the WT tomogram). Scale bar, 200 nm. For full tomographic reconstructions and segmentations, see Supplementary Videos 1–3. **(h)** Representative plaque assay images of the parental (DiCreΔ*Ku80*) and *loxPpckmt-Halo* strains after 7 days ± 50 nM rapamicin (n = 3). **(i)** Quantification of invasion efficiency after 72 h ± 50 nM rapamicin. Data are normalised to the parental control (DiCreΔ*Ku80*). Bars show mean ± SD; white dots represent means of each biological replicate (n = 3). Statistical significance was determined by two-tailed Student’s t-test; *** p < 0.001. **(j)** Quantification of gliding speed. Box-and-whisker plots show the median and interquartile range; dots represent individual parasites pooled across three biological replicates. Statistical significance was determined by two-tailed Student’s t-test; *** p < 0.001.

To characterise the role of PCKMT in the lytic cycle, we introduced loxP sites flanking the *pckmt* locus in a strain expressing DiCre recombinase (Supplemenatary Fig. 1c-e)^17,18^. Time-course quantification revealed that by 48 hours post-induction, over 80% of the population had lost detectable PCKMT signal, reaching near-complete 100% knockout by 72 hours (Fig. 1c, Supplemenatary Fig. 1f).

Given its identification as a CGP-associated factor, we next asked whether PCKMT depletion disrupts the localisation of apical proteins that depend on the PCRs for recruitment. Strikingly, loss of PCKMT resulted in a complete loss of FRM1 from the PCRs (Fig. 1d), establishing PCKMT as essential for anchoring this key actin nucleator at the apex. Conversely, FRM1 depletion did not affect PCKMT localisation, indicating a unidirectional dependency (Supplemenatary Fig. 1g). DAP1, another conoid-associated factor that is dispensable for parasite survival^3^, also exhibited partial mislocalisation upon PCKMT depletion (Supplemenatary Fig. 1h, i), whereas ICAP16 remained correctly positioned (Supplemenatary Fig. 2a). Notably, ICAP16 was previously reported to contribute moderately to host cell invasion^19^. These findings indicate that PCKMT selectively stabilises a subset of apical complex proteins, with FRM1 being its most prominent effector.

**Fig. 2:**
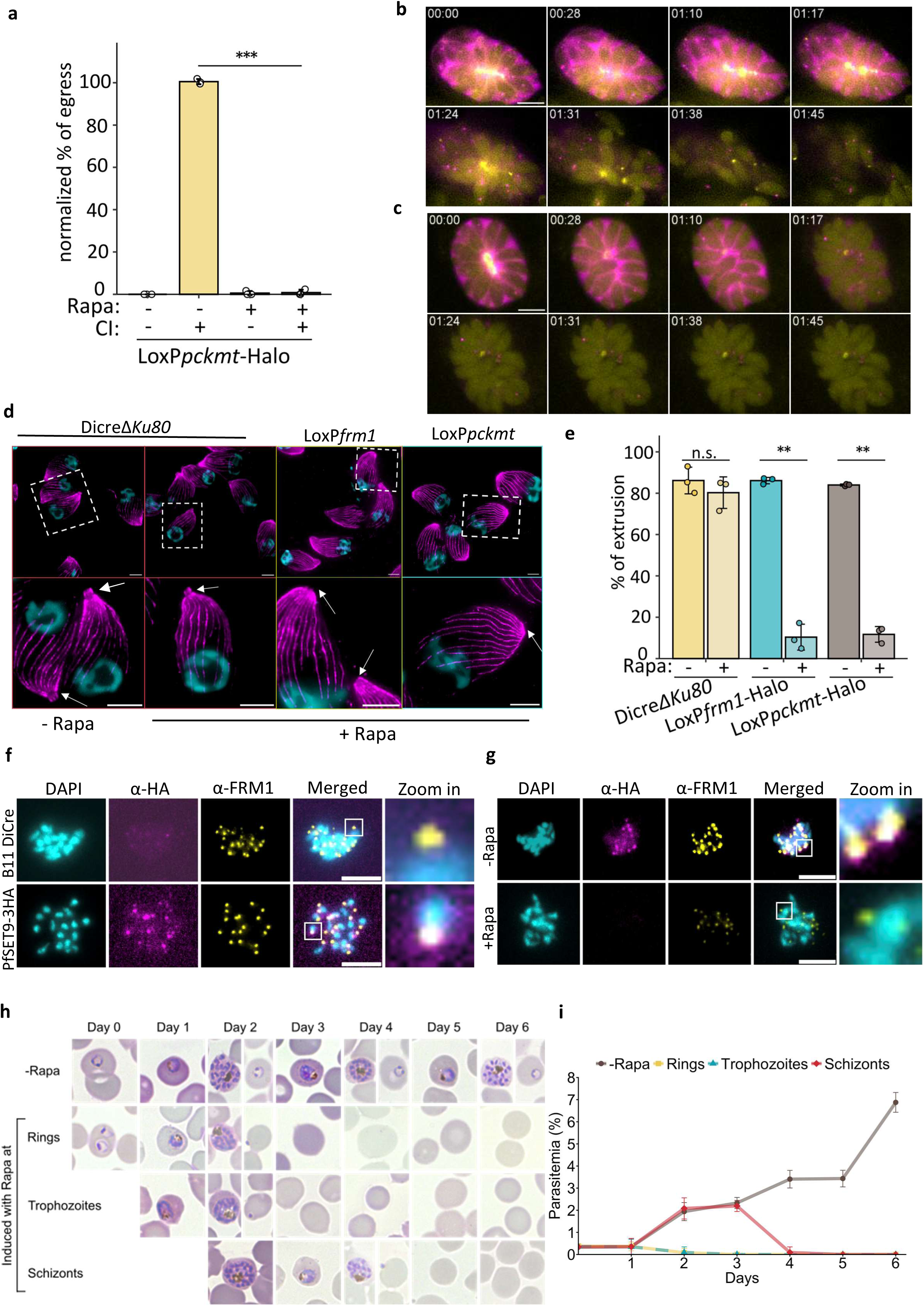
Conserved apical methyltransferase function: PCKMT and PfSET9 are required for parasite propagation. **(a)** Quantification of *T. gondii* egress efficiency using a fixed-egress assay. Parasites were pre-induced with or without rapamycin (Rapa) for 72 h, and egress was triggered with 4 µM calcium ionophore A23187 (CI) for 5 min. Egress efficiency was normalised to the DiCreΔ*Ku80* control. Bars show mean ± SD; white dots indicate biological replicates (n = 3). Statistical significance was assessed by two-tailed Student’s t-test; *** p < 0.001. **(b–c)** Representative stills from live egress imaging (Movie S4). Parasites expressing Chromobody-emerald (CbEm; yellow, F-actin) and SAG1ΔGPI-dsRed (magenta, parasitophorous vacuole) were stimulated with 4 µM CI. **(b)** Non-induced parasites show actin remodelling and egress. **(c)** PCKMT-depleted parasites fail to initiate motility despite vacuole rupture. Scale bar, 5 µm. **(d)** Expansion microscopy of conoid extrusion following ∼10 min CI stimulation. White arrows mark extruded conoids. Scale bar, 5 µm. **(e)** Quantification of conoid extrusion. ≥100 parasites were counted per condition. Bars show mean ± SD; dots represent biological replicates (n = 3). Two-tailed Student’s t-test; ** p < 0.01; n.s., non-significant. **(f)** Immunofluorescence of *P. falciparum* schizonts showing apical PfSET9 (the PCKMT orthologue) co-localising with FRM1 at the apical complex of segmented merozoites. Scale bar, 5 µm. **(g)** PfSET9-depleted schizonts (48 h post-induction with 100 nM Rapa) show reduced PfSET9 and weaker FRM1 signals. Scale bar, 5 µm. **(h)** Representative Giemsa-stained smears of synchronous cultures induced at ring, trophozoite, or schizont stages, showing severe invasion defects after PfSET9 depletion. **(i)** Quantification of parasite growth over 6 days reveals a pronounced growth defect following PfSET9 depletion.

To exclude potential conoid structural defects similar to those reported in CGP-depleted parasites^3^, we next examined whether loss of PCKMT affected the organisation of other preconoidal components. CGP localisation was not affected by PCKMT depletion, and likewise, Pcr4 remained correctly positioned at the preconoidal rings (Fig. 1e, f), indicating that PCKMT is not required for maintaining overall preconoidal integrity. To further rule out subtle structural perturbations, we performed cryo-electron tomography (cryo-ET), which revealed a morphologically intact conoid and associated apical complex in mature PCKMT-KO parasites (Fig. 1g; Supplemenatary Fig. 2b–e; Movies S1–S3).

Having established that loss of PCKMT selectively affects specific apical factors without grossly altering conoid architecture, we next assessed its functional relevance during the lytic cycle. Plaque assays revealed a complete loss of parasite growth after one week of induction with 50 nM rapamycin (Fig. 1h). Replication assays confirmed that intracellular division proceeds normally in the absence of PCKMT (Supplemenatary Fig. 3a), indicating that the loss of PCKMT primarily affects motility-associated processes such as invasion, egress, or gliding. Indeed, parasites lacking PCKMT exhibited a severe invasion defect, comparable to knockouts of core motility factors, including actin or MyoA^20^ (Fig. 1i). Gliding motility was also profoundly impaired: only a small fraction of parasites initiated motility, and those that did moved at reduced speed (Fig. 1j; Supplemenatary Fig. 3b; Movie S4).

**Fig. 3:**
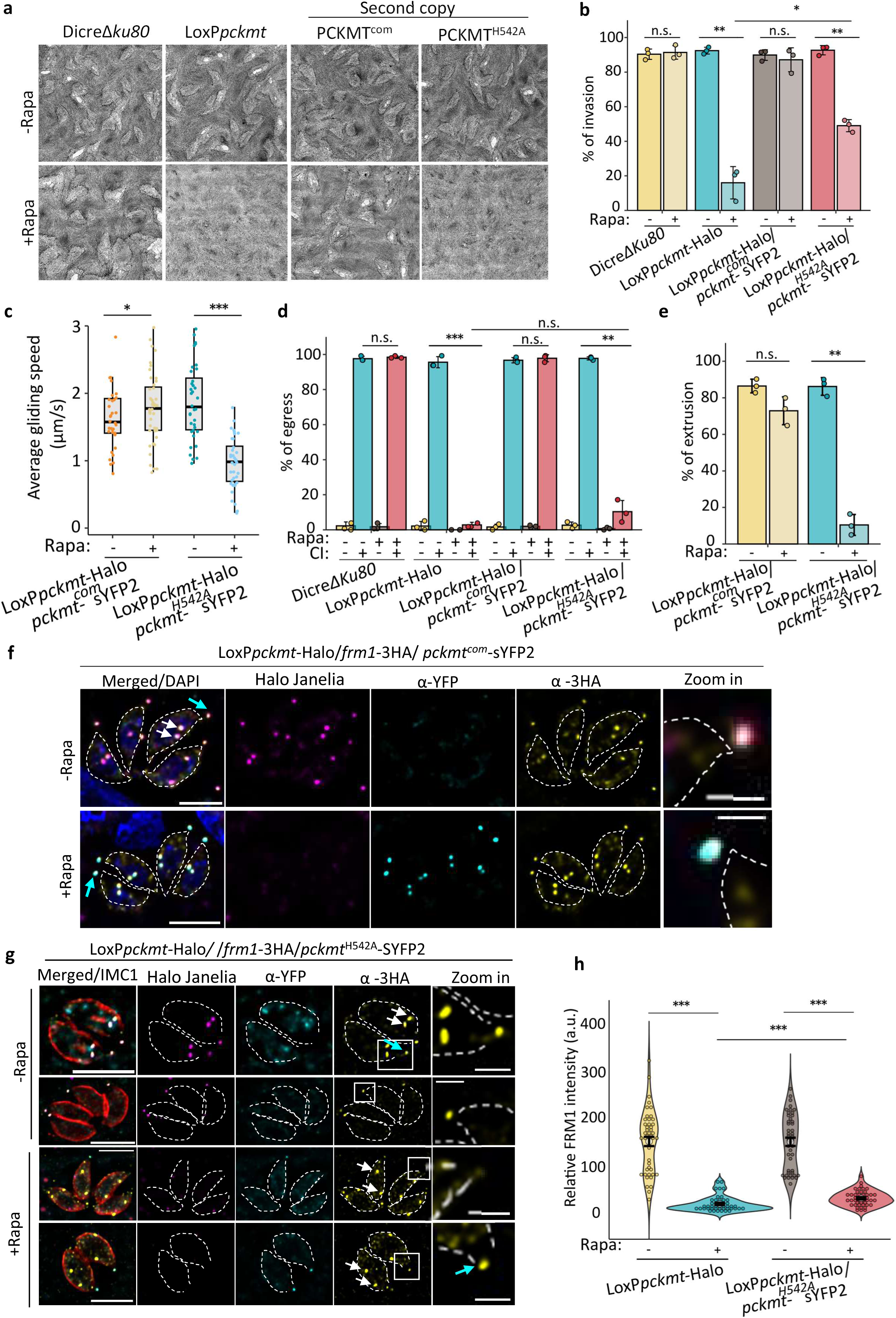
PCKMT catalytic activity is essential for FRM1 anchoring and motility initiation. **(a)** Representative plaque assays of *T. gondii* DiCreΔ*Ku80* (parental), loxPpckmt-Halo, loxPpckmt-Halo/PCKMT^com^ (complemented), and loxPpckmt-Halo/PCKMT^H542A^ (catalytic-dead) strains after ∼7 days ± 50 nM rapamycin (Rapa). Scale bar, 5 mm. Details of mutant construction are shown in Supplemenatary Fig. S5. **(b)** Quantification of invasion efficiency in the indicated strains post induction ± Rapa for 72 h. Bars show mean ± SD; white dots represent biological replicates (n = 3). **(c)** Quantification of gliding motility speed in PCKMT^com^ and PCKMT^H542A^ parasites ± Rapa. Box-and-whisker plots show median and interquartile range; dots represent individual parasites pooled from three biological replicates. **(d)** Quantification of egress efficiency in PCKMT^com^ and PCKMT^H542A^ parasites pre-induced with or without Rapa. Egress was triggered with 4 µM calcium ionophore A23187 (CI) for 5 min. ≥100 vacuoles were scored per condition. Bars show mean ± SD; white dots represent biological replicates (n = 3). **(e)** Quantification of conoid extrusion in PCKMT^com^ and PCKMT^H542A^ parasites ± Rapa after CI stimulation for ∼10 min. ≥100 parasites were analysed per condition. Bars show mean ± SD; white dots represent biological replicates (n = 3). **(f)** Immunofluorescence showing that PCKMT^com^ expression is maintained after rapamycin-induced depletion of endogenous PCKMT, with FRM1 retaining normal localisation at the conoid. Cyan arrows indicate mature parasites; white arrows mark daughter cells. Scale bar, 5 µm (merged view), 1 µm (zoom). **(g)** Representative image showing reduced FRM1 localisation in PCKMT^H542A^ parasites ± Rapa. The white arrow indicates conoidal PCKMT^H542A^ signal. Three biological replicates were performed. **(h)** Quantification of relative FRM1 intensity in the indicated strains. Each dot represents the mean FRM1 intensity per vacuole (∼15 vacuoles per biological replicate; n = 3). Statistical significance for all quantifications in this figure was assessed by a two-tailed Student’s *t*-test; * *= p* < 0.05, ** *= p* < 0.01, *** *= p* < 0.001, n.s.: non-significant.

Consistent with these defects, PCKMT-depleted parasites also failed to egress from host cells even when stimulated with calcium ionophore (Fig. 2a). To dissect this phenotype in more detail, we analysed actin dynamics during induced egress using parasites expressing chromobody-emerald (CbEm) to visualise F-actin and SAG1ΔGPI-dsRed marker^2^ to visualise the parasitophorous vacuole space. In non-induced parasites, actin filaments depolymerised upon stimulation, the vacuole ruptured, and F-actin relocalised to the parasite basal end, enabling effective egress (Fig. 2b; Movie S5). In contrast, PCKMT-depleted parasites exhibited a pronounced egress block: although the parasitophorous vacuole ruptured, F-actin failed to accumulate at the posterior pole, indicating a failure in motility initiation reminiscent of CGP-deficient parasites^2^ (Fig. 2c; Movie S5).

Since motility initiation is tightly linked to conoid protrusion^21^, we used expansion microscopy to compare wild-type, *pckmt*-induced knockout (iKO), and *frm1*-iKO parasites. In both PCKMT- and FRM1-depleted parasites, conoid extrusion was significantly impaired compared to controls (Fig. 2d, e), suggesting that PCKMT acts upstream of F-actin polymerisation.

To explore whether the function of PCKMT might be conserved across apicomplexan parasites, we next examined its *Plasmodium falciparum* orthologue, annotated as *set9*. This gene was previously detected in transcriptional signatures of sexually committed parasites^22^, but its localisation and function have not been described so far. We generated a conditional mutant by inserting a loxP intron into the *set9* locus in a DiCre background^23,24^ (Supplemenatary Fig. 3c–e). Immunofluorescence analysis revealed that SET9 localises as a distinct apical dot, in close proximity with PfFRM1, in segmented merozoites during the asexual blood stage (Fig. 2f). Upon SET9 depletion, FRM1 signal appeared reduced or partially mislocalised compared to controls, suggesting that proper FRM1 positioning depends, at least in part, on SET9 activity and indicating a conserved functional link between PCKMT/SET9 and FRM1-like factors in regulating apical assembly and invasion (Fig. 2g). Following rapamycin-induced excision at different developmental stages (Supplemenatary Fig. 3f), parasites failed to transition from schizont to ring stages and exhibited a pronounced growth arrest (Fig. 2h, i), consistent with an invasion defect reminiscent of the phenotype observed in *T. gondii* PCKMT knockouts.

Together, these data identify PCKMT as an apical lysine methyltransferase that is essential for conoid-dependent motility in *T. gondii* and point to an evolutionarily conserved role for its *Plasmodium* orthologue SET9 in invasion-associated processes across apicomplexans. To explore how PCKMT exerts its function, we tested whether its methyltransferase activity, mediated by a conserved SET domain, is required for FRM1 recruitment and motility initiation.

### Catalytic activity of PCKMT is required for anchoring FRM1 and triggering parasite motility

The catalytic histidine (H542) lies within the highly conserved HxC motif, a hallmark of active SET-domain methyltransferases. Sequence alignment of PCKMT with AKMT, other apicomplexan SET-domain enzymes, and the human cytoskeletal methyltransferase SMYD2^25^ confirmed conservation of key catalytic residues across species (Supplemenatary Fig. 4a), supporting its likely enzymatic activity. This residue corresponds to H447 in AKMT, which was previously shown to be essential for methyltransferase activity when mutated to valine^15^. Based on this conservation, we introduced a point mutation substituting the catalytic histidine with alanine (H542A) in the PCKMT SET domain to disrupt catalytic activity.

**Fig. 4:**
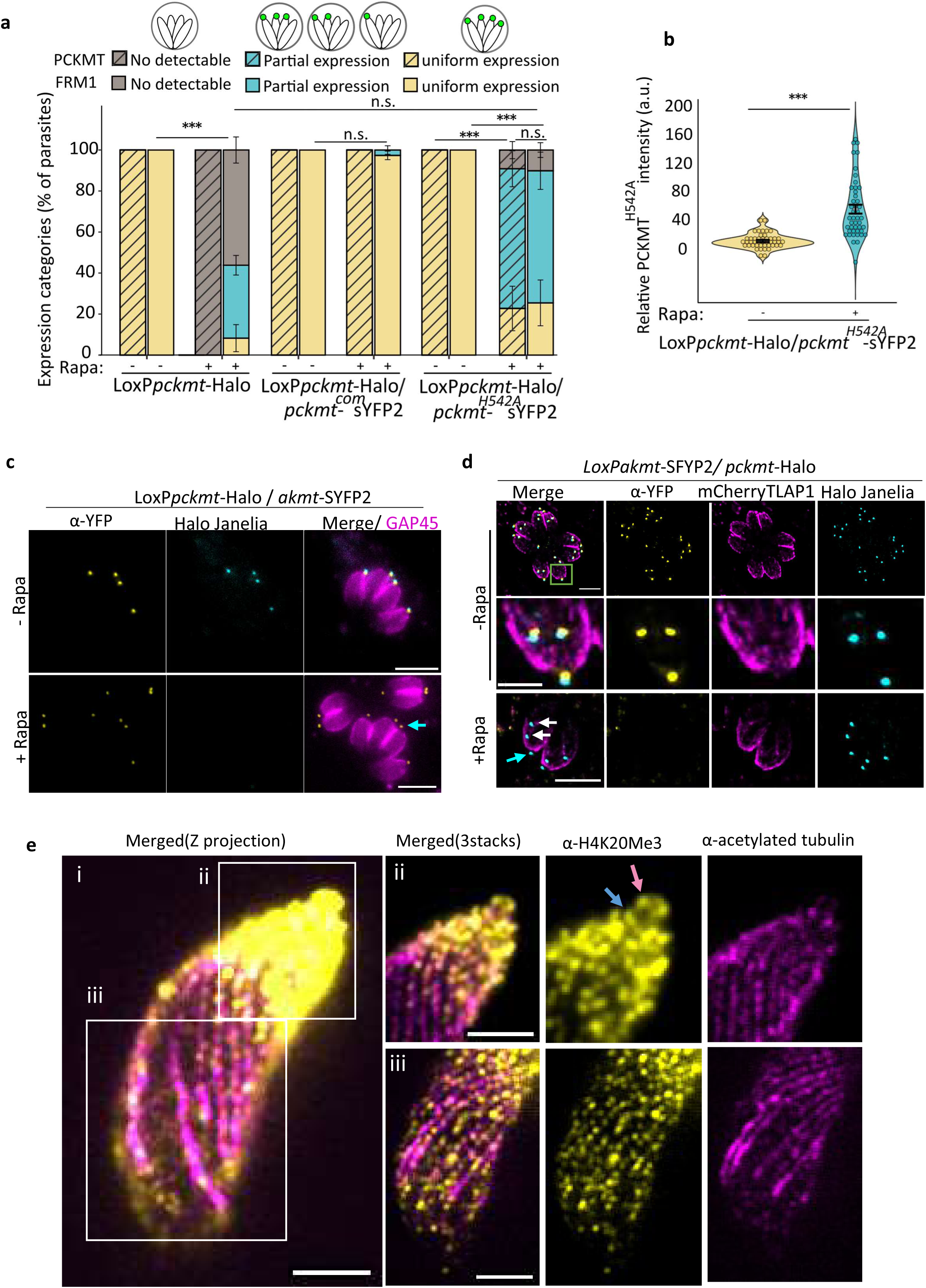
Quantitative conoid localisation of PCKMT^H542A^ and FRM1, reciprocal independence of AKMT and PCKMT, and STED mapping of apical methylation. **(a)** Quantification of PCKMT^H542A^ and FRM1 expression at the conoid. Parasite vacuoles were classified into three categories: (1) no detectable expression within the vacuole, (2) partial expression, where only a subset of parasites displayed detectable conoidal signal, and (3) uniform expression, where all parasites within the vacuole showed conoidal localisation. Proteins with even faint signals were scored as expressed regardless of intensity. ≥100 vacuoles were analysed per condition across three biological replicates. Bars show mean ± SD. **(b)** Quantification of relative PCKMT^H542A^ intensity at the conoid in the loxPpckmt-Halo/pckmt^H542A^-SYFP2 strain. Each dot represents the mean conoid intensity per vacuole (∼15 vacuoles per biological replicate; n = 3). **(c)** AKMT localisation remains unaltered upon PCKMT depletion. Cyan arrow indicates AKMT at the mature conoid. Scale bar, 5 µm. **(d)** PCKMT localisation remains unaltered upon AKMT depletion. Cyan arrow indicates PCKMT at the mature conoid. White arrow: daughter cell conoids Scale bar, 5 µm. Zoom in scale bar, 2 µm. **(e)** STED microscopy of extracellular *T. gondii* tachyzoites. (i) Maximum-intensity z-projection showing overall methylation pattern. (ii) Merge of three optical sections highlighting apical structures, with strong methylation signals at the preconoidal rings (pink arrow) and apical polar ring (blue arrow). (iii) Methylation puncta align along cortical microtubules. Scale bar, 1 µm. Statistical significance (a–b) was assessed by a two-tailed Student’s *t*-test; * *= p* < 0.05, ** *= p* < 0.01, *** *= p* < 0.001, n.s.: non-significant.

To evaluate the functional impact of this substitution, we generated complementation lines expressing a second copy of either wild-type PCKMT (PCKMT^com^) or the mutant version (PCKMT^H542A^) in the inducible knockout background (Supplemenatary Fig. S4b, S5).

Complementation of the PCKMT-inducible knockout (iKO) strain with PCKMT^com^ fully restored parasite growth in plaque assays, while the PCKMT^H542A^ mutant failed to rescue growth (Fig. 3a). Further analysis revealed that PCKMT^com^ rescued all steps of the lytic cycle, including invasion, gliding, egress, and conoid extrusion (Fig. 3b-e; Supplemenatary Fig. 6a, b). In contrast, the PCKMT^H542A^ line exhibited a partial rescue: invasion and gliding were moderately improved compared to the knockout, but egress remained severely impaired (Fig. 3a-d; Supplemenatary Fig. 6a; Movie S6). Conoid extrusion was also defective in the mutant line, indicating that catalytic activity is required for this critical early step in motility initiation (Fig. 3e, Supplemenatary Fig. 6b, c).

**Fig. 5:**
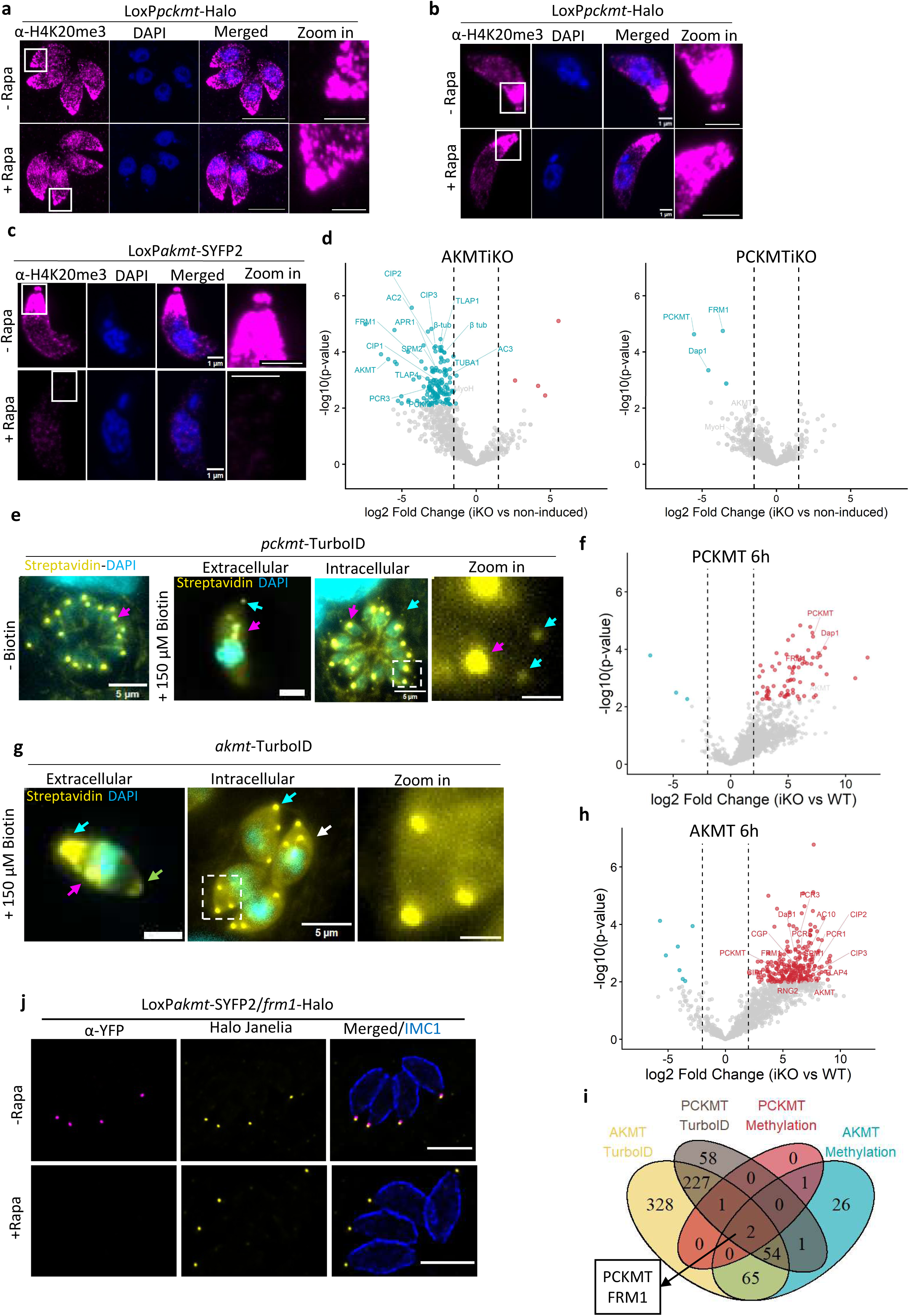
Apical lysine methyltransferases establish a conoid methylation network controlling FRM1 localisation in *T. gondii*. **(a–c)** Immunofluorescence analysis of methylation using anti-H4K20me3 antibody in parasites induced ± rapamycin (Rapa). (a) Intracellular *loxPpckmt*-*Halo* parasites. (b) Extracellular *loxPpckmt-Halo* parasites. (c) Extracellular *loxPakmt-YFP* parasites. Comparable methylation patterns were observed with α-H3K36me3, α-Kme1/2, and α-Pan-Kme3 antibodies (Supplemenatary Fig. S7c,d). Scale bars, 5 µm. **(d)** Volcano plots showing differential methyl-peptide enrichment between induced and non-induced parasites for *loxPpckmt-Halo* and *loxPakmt-YFP* strains. **(e)** Representative images of *pckmt*–*TurboID* parasites stained with streptavidin (yellow) and DAPI (cyan). In the absence of biotin, a strong streptavidin signal is detected at the apicoplast (magenta arrows). After 6 h induction with 150 µM biotin, additional puncta appear at the conoid (white arrows). Insets show conoidal localisation. Scale bars, 5 µm. **(f)** Volcano plots from TurboID proximity-labelling experiments for PCKMT after 6 h biotin induction. **(g)** Representative images of AKMT–TurboID parasites stained with streptavidin (yellow) and DAPI (cyan). Following 6 h biotin induction, signal is detected both apically (white arrows) and at the basal complex (green arrows) besides the expected localisation at the apicoplast (magenta arrow). Insets show apical puncta with mature (cyan arrow) and daughter cells (white arrows). Scale bars, 5 µm. **(h)** Volcano plots from TurboID proximity-labelling experiments for AKMT after 6 h biotin induction. **(i)** Venn diagram summarising overlap between methyl-enrichment and TurboID datasets, highlighting PCKMT and FRM1 as the only proteins consistently identified across all analyses. **(j)** Endogenous tagging of FRM1 in conditional *loxPakmt-YFP* parasites shows that FRM1 is not lost from the conoid upon AKMT depletion, confirming a unidirectional dependency. Scale bars, 5 µm.

**Fig. 6:**
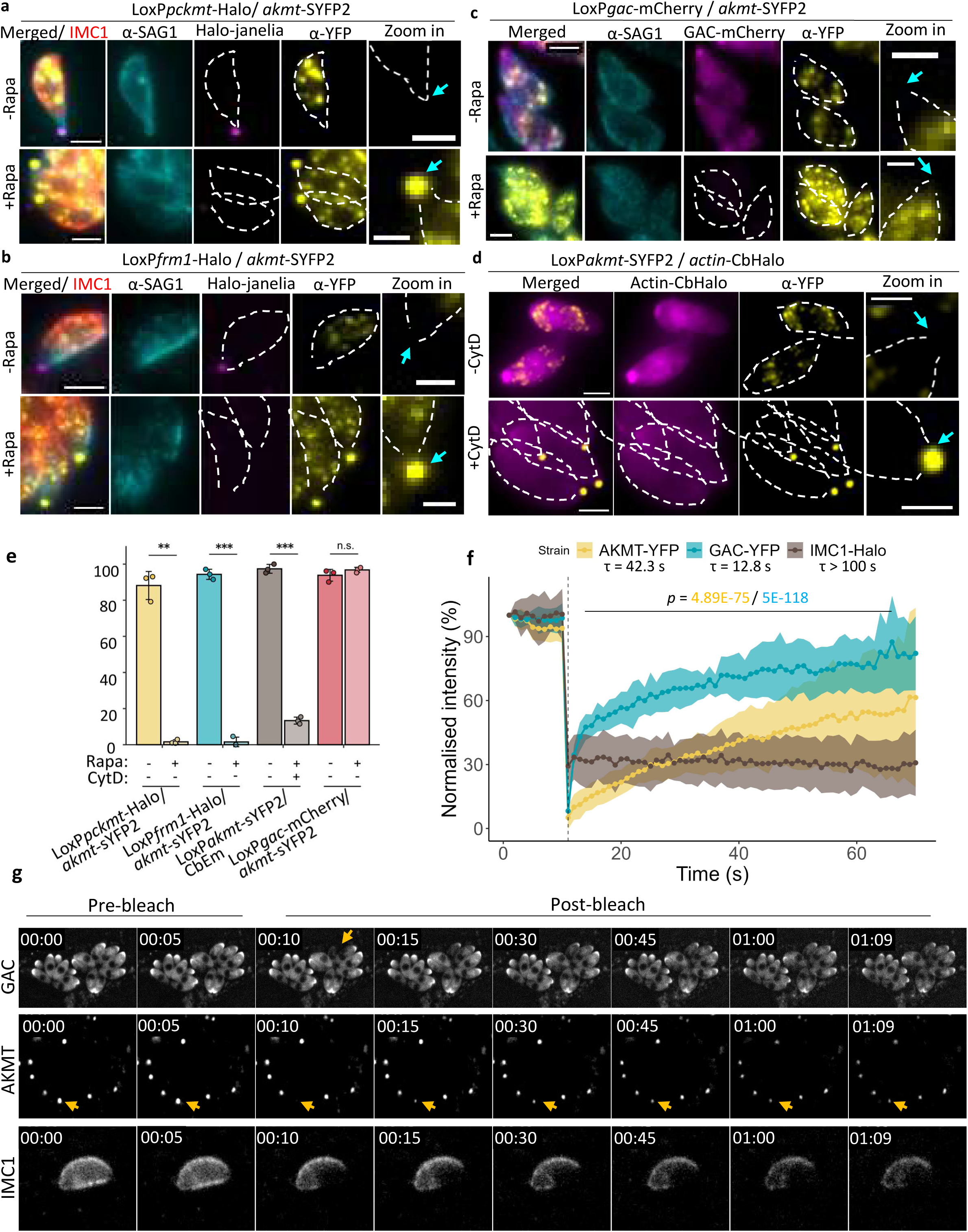
Dynamic relocalisation of AKMT requires PCKMT and actin polymerisation. **(a–b)** Immunofluorescence analysis of AKMT dynamics during calcium-triggered egress in *loxPpckmt-Halo* (a) and *loxPfrm1-Halo* (b) parasites induced ± rapamycin (Rapa) for 54 h. Egress was stimulated with 4 µM calcium ionophore A23187 (CI) for 5 min. Anti-SAG1 (cyan, non-permeabilising) marks parasites that have ruptured the parasitophorous vacuole membrane (PVM) but not completed egress. Cyan arrows indicate localization of the conoid; scale bars, 2 µm (merged) and 1 µm (zoom). **(c)** AKMT translocation occurs independently of GAC. *loxPgac-Halo* parasites were induced ± Rapa for ≥ 54 h and stimulated with 4 µM CI for 5 min. Anti-SAG1 (cyan) marks lysed vacuoles. Cyan arrows highlight the absence of AKMT at the conoid. Scale bars, 2 µm (merged) and 1 µm (zoom). **(d)** AKMT relocalisation depends on actin polymerisation. Parasites expressing CbHalo-labelled F-actin were treated with 2 µM cytochalasin D (CytD) for 1 h before egress induction with 4 µM CI for 5 min. In CytD-treated parasites, AKMT remains apical (cyan arrows). Scale bars, 2 µm (merged) and 1 µm (zoom). **(e)** Quantification of AKMT translocation from panels (a–d). In the +Rapa condition, only parasites lacking the corresponding floxed protein (PCKMT, FRM1, or GAC) were considered. Bars show mean ± SD; dots represent the mean of each biological replicate (n = 3). Statistical significance was assessed by two-tailed Student’s t-test (**p < 0.01, ***p < 0.001, n.s. = not significant). **(f)** Fluorescence-recovery-after-photobleaching (FRAP) analysis of apical proteins. Parasites expressing the indicated fluorescently tagged constructs were imaged for 10 s before bleaching at the apical tip, followed by 60 s recovery. The dotted vertical line marks the bleaching time point. Normalised fluorescence intensity (mean ± SD) is plotted over time (n = 3 biological replicates). Recovery rates (τ) were obtained by exponential fitting. A two-way repeated-measures ANOVA compared recovery dynamics to the non-recovering control IMC1-Halo; p-values are colour-coded as in the legend. **(g)** Montage of selected frames from Movie S6 illustrating the rapid fluorescence recovery quantified in (f). Orange arrows denote the bleached region.

We next examined how the catalytic mutation in PCKMT affects FRM1 recruitment to the conoid. To do this, we performed immunofluorescence assays in parasites expressing second-copy complementation lines. Unexpectedly, expression of the second copy, either PCKMT^WT^ or PCKMT^H542A^, was only robustly detected upon depletion of the endogenous copy, rather than in parallel with it as anticipated (Fig. 3f, g). This may reflect integration site effects at the *uprt* locus or a very tight regulation of PCKMT expression.

In the complemented strain, FRM1 remained stably anchored at the conoid, whereas in the PCKMT^H542A^ mutant, FRM1 was largely absent from mature tachyzoites (Fig. 3g) but still detectable in developing daughter buds, consistent with previous observations that FRM1 is recruited early to nascent conoids during cell division^3^ (white arrows). This pattern suggests that the retention of FRM1, and potentially other conoidal proteins, within the mature conoid requires a post-budding maturation step. Quantification of FRM1 fluorescence intensity confirmed a strong reduction at mature conoids in the mutant strain, nearly comparable to the full PCKMT knockout (Fig. 3h). Closer inspection revealed heterogeneous FRM1 expression within vacuoles: in the catalytic-inactive mutant, approximately 64% of vacuoles contained a mixed population of parasites with or without FRM1 signal, whereas only 25% displayed uniform FRM1 localisation at mature conoids (Fig. 4a). These results indicate that the catalytic activity of PCKMT is required for stable FRM1 recruitment and retention at the mature conoid, whereas its initial incorporation during daughter conoid formation can occur independently of catalysis, consistent with previous observations that several PCR components transiently associate with budding conoids during division before disengaging upon maturation^3^.

Interestingly, not all vacuoles expressed detectable levels of the second-copy PCKMT^H542A^ protein, raising the possibility that an active SET domain may be required for the proper recruitment or stabilisation of PCKMT itself at the conoid (Fig. 4a, b). In fact, AKMT^H447V^ also failed to localise fully at the conoid^15^. This suggests a potential feedback mechanism in which enzymatic activity reinforces localisation or retention of these methyltransferases at their site.

Taken together, these results demonstrate that PCKMT’s catalytic activity is essential for anchoring FRM1 at the conoid and initiating parasite motility. The inability of the H542A mutant to restore growth, conoid extrusion, or egress confirms that enzymatic activity, rather than mere structural presence, is required for PCKMT function. The reduced localisation of both FRM1 and the inactive PCKMT variant further suggests a feedback mechanism in which catalytic activity stabilises PCKMT and its partners at the conoid.

### PCKMT and AKMT localise independently at the apical complex

PCKMT is not the only methyltransferase localised at the conoid. To investigate potential functional interplay with the apical methyltransferase AKMT, which relocalises to the apical complex during motility activation, we examined AKMT dynamics and the reciprocal localisation of both enzymes in their respective knockout backgrounds.

AKMT is a SET-domain methyltransferase shown to regulate gliding motility^15^ by promoting the recruitment of GAC, which links F-actin to the parasite surface adhesins^9^. Immunofluorescence analysis revealed that PCKMT depletion did not affect AKMT localisation at the conoid in intracellular parasites (Fig. 4c), nor did it impact GAC recruitment (Supplemenatary Fig. 6d). Conversely, AKMT knockout did not alter the localisation of PCKMT, which remained at the preconoidal rings (Fig. 4d). These results indicate that PCKMT and AKMT localise independently and may function in parallel to prepare the conoid and associated cytoskeletal components for motility activation.

### Distinct Methylation Networks Governed by PCKMT and AKMT

Next, to further dissect the functional distinction between PCKMT and AKMT, we analysed global methylation patterns using an antibody specific for trimethylated lysine 20 of histone H4 (α-H4K20me3)^26^. Interestingly, this antibody revealed a broad methylation profile concentrated at the apical complex and extending along the subpellicular microtubules (Fig. 4e, Supplemenatary Fig. 7a).

**Fig. 7:**
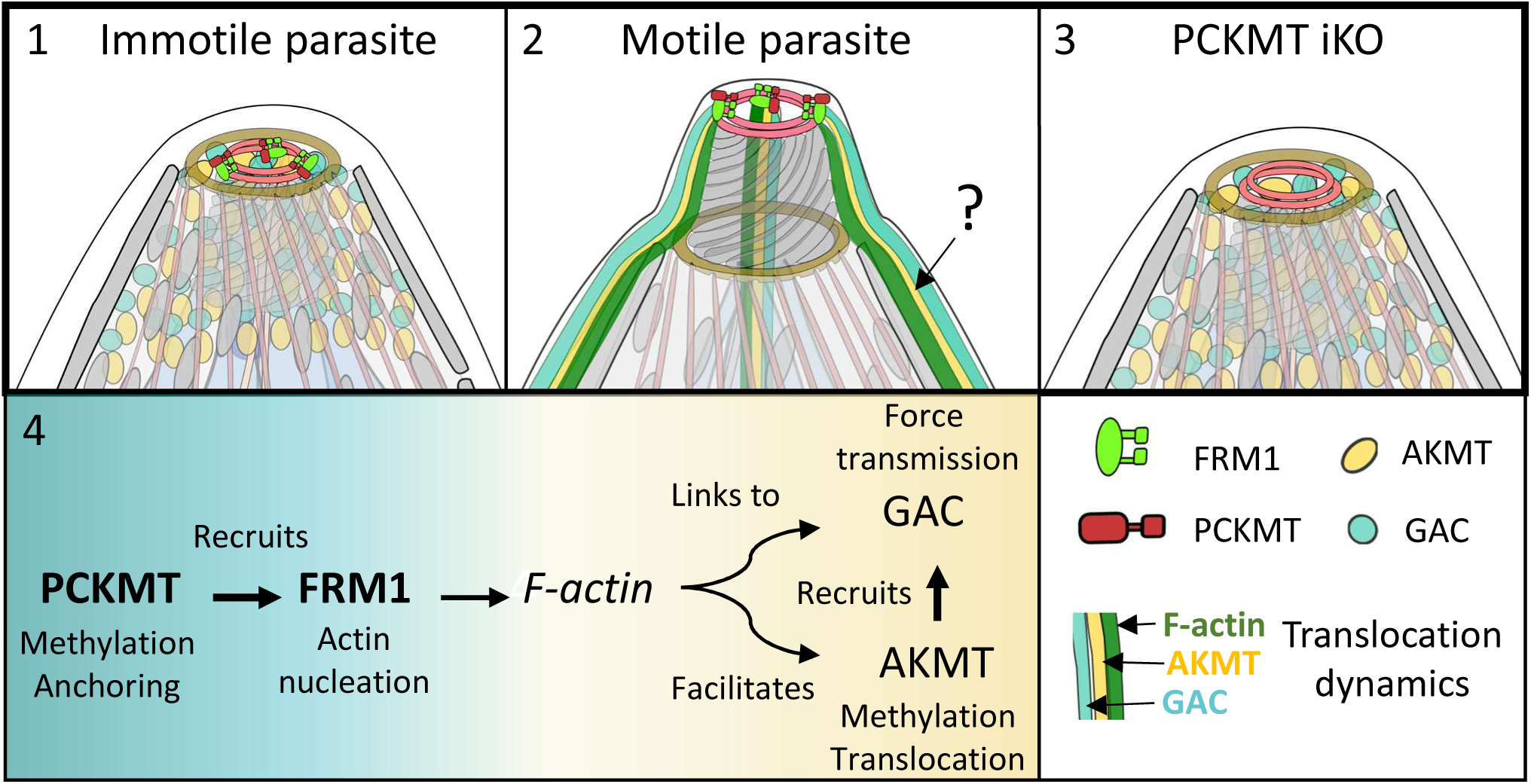
Coordinated methylation logic driving motility initiation in apicomplexan parasites. **(1)** In non-motile parasites, PCKMT, AKMT, GAC, and FRM1 are pre-assembled at the apical complex, poised for rapid activation. **(2)** Upon stimulation, PCKMT enables FRM1-dependent actin nucleation at the conoid, promoting conoid protrusion and the apical-to-basal translocation of AKMT and GAC, which couple F-actin to surface adhesins to drive motility. The exact route of AKMT movement (question mark) remains unresolved—whether it traverses the sub-pellicular space between the plasma membrane (PM) and the inner membrane complex (IMC) or diffuses through the cytosol is unknown. **(3)** In the absence of PCKMT, FRM1 fails to localise to the conoid, actin assembly does not occur, and AKMT/GAC translocation is blocked, preventing motility initiation. **(4)** The lower panel summarises the temporal sequence of activation, PCKMT → FRM1 → actin → GAC/AKMT, illustrating the cascade that links methylation to cytoskeletal activation and force transmission.

In intracellular PCKMT-depleted parasites, we observed no detectable change in H4K20me3 signal compared to non-induced parasites (Fig. 5a), suggesting that PCKMT does not contribute to global methylation patterns under these conditions. In extracellular parasites, the signal was more elevated than in intracellular parasites, suggesting an increase in methylation after motility initiation, but again unchanged in the absence of PCKMT (Fig. 5b).

In contrast, depletion of AKMT resulted in a striking loss of methylation signal in extracellular parasites (Fig. 5c). This confirms AKMT as the dominant contributor to methylation detectable by these antibodies during the lytic cycle as previously described^9^. To address the possibility that PCKMT acts on a restricted subset of targets not detectable by population-level immunofluorescence, we generated a double knockout strain lacking both AKMT and PCKMT, hereafter referred to as the Double Apical MethylTransferase inducible KnockOut (DAMTiKO). However, this mutant displayed methylation levels indistinguishable from the AKMT single knockout (Supplemenatary Fig. 7b), further supporting a limited role for PCKMT in bulk lysine methylation.

Comparable methylation patterns were observed using alternative antibodies, including α-H3K36me3, α-Kme1/2, and α-Pan-Kme3, in both intracellular and extracellular parasites (Supplemenatary Fig. 7c, d). Although these antibodies might recognise partially overlapping methylation marks and can display sequence bias^27^, their consistent apical labelling across different reagents strongly supports that the observed methylation pattern reflects genuine, biologically relevant modifications rather than antibody cross-reactivity. Consistent with this, western blot analysis confirmed a corresponding reduction in methylation levels upon enzyme depletion (Supplemenatary Fig. 8a).

To explore whether PCKMT targets a restricted subset of substrates, we performed α-H4K20me3– mediated methylated protein enrichment (Fig. 5d; Supplemenatary Fig. 9a, b) and TurboID-based proximity labelling (Fig. 5e-h; Supplemenatary Fig. 9c-f) in both wild-type and KO backgrounds. Comparative analysis of these proteomes revealed largely distinct sets of interactors for PCKMT and AKMT, with only FRM1 and PCKMT emerging as a consistent hit in all datasets (Fig. 5i). To further characterise these datasets, we performed Gene Ontology (GO) enrichment analysis of the proteins reduced upon AKMT or PCKMT depletion (Supplemenatary Fig. 9b). AKMT-associated proteins were significantly enriched for cytoskeletal organisation and motility-related processes, including the myosin complex, actin cytoskeleton, and vesicle organisation, consistent with its broad role in apical and pellicular remodelling. By contrast, the limited number of proteins affected in the PCKMT methylome yielded no meaningful GO enrichment, in line with its narrow substrate specificity. Similar trends were observed in the TurboID datasets (Supplemenatary Fig. 9e), reinforcing the functional partitioning between AKMT and PCKMT revealed by the proteomic analyses.

To validate whether FRM1 localisation depends on AKMT, we endogenously tagged FRM1 in the AKMT-floxed line and monitored its distribution upon depletion. In contrast to the loss of FRM1 from the conoid observed in PCKMT-depleted parasites (Fig. 1d), FRM1 localisation remained unaffected following AKMT depletion (Fig. 5i), confirming that AKMT is not required for FRM1 anchoring at the apical tip.

We also tagged several additional candidates that appeared in both the AKMT and PCKMT TurboID datasets and localised to the conoid of mature or developing parasites, including the recently described apical polar ring scaffold ASAF1^3^. None of these proteins showed altered localisation following PCKMT depletion (Supplemenatary Fig. 10a-e).

Together, these findings demonstrate that PCKMT and AKMT define distinct yet complementary methylation networks at the apical complex. PCKMT acts on a narrow subset of substrates, most notably anchoring FRM1 to the preconoidal rings, whereas AKMT methylates a broader range of cytoskeletal targets at the conoid and pellicle. This division of labour explains the contrasting methylation patterns observed by immunofluorescence and underscores the robustness of our combined proteomic and imaging-based analyses.

### PCKMT links actin assembly to AKMT mobilisation at the conoid

In the current model of motility initiation, signalling cues trigger conoid protrusion^21,28^, followed closely by the disengagement of AKMT^15^ from the apex and the translocation of GAC^9^ together with F-actin to the parasite’s basal end to generate the force needed for gliding. To position PCKMT within this cascade, we analysed AKMT dynamics during egress induction. To capture the onset of AKMT translocation, parasites were stimulated with a calcium ionophore before fixation, and lysed parasitophorous vacuoles were stained with anti-SAG1 under non-permeabilising conditions to identify those that had initiated egress, including in the induced knockout lines that are severely impaired in motility. In wild-type parasites, calcium ionophore stimulation triggers AKMT disengagement from the conoid and relocalisation to the posterior end. In contrast, PCKMT-depleted parasites retained AKMT at the conoid in most cells, with quantification revealing a near-complete block in translocation (Fig. 6a; Supplemenatary Fig. 10f).

We next examined whether the conoid-associated formin FRM1^8^ acts upstream of AKMT translocation. FRM1 depletion phenocopied the PCKMT knockout, with AKMT failing to relocalise upon ionophore stimulation (Fig. 6a; Supplemenatary Fig. 10g). These data position FRM1 as a key factor upstream of AKMT mobilisation and suggest that PCKMT may promote local actin polymerisation by recruiting or activating FRM1 at the conoid. Finally, to refine pathway order, we analysed AKMT localisation in parasites lacking GAC. Contrary to earlier reports^9^, AKMT readily disengaged from the conoid in GAC knockout parasites, although it appeared more diffusely distributed within the cytosol (Fig. 6b; Supplemenatary Fig. 10h).

To test whether actin polymerisation is required for AKMT relocalisation, we treated wild-type parasites with Cytochalasin D (CytD). Following ionophore induction, AKMT remained concentrated at the conoid (Fig. 6c; Supplemenatary Fig. 10i), consistent with previous reports^9,15^. Notably, the cytosolic AKMT subpopulation observed in the absence of PCKMT or FRM1, was frequently diminished or absent under CytD treatment, suggesting that actin dynamics facilitate its redistribution within the cytosol, even when conoid disengagement is blocked. The sporadic presence of diffuse AKMT signal further implies that additional F-actin nucleation centres, such as those formed by Formin 2 (FRM2) near the Golgi–apicoplast region^17^, might contribute to this residual mobility.

We quantified these dynamics across all conditions in Fig. 6d, including only parasites deficient for the floxed genes analysed above or treated with CytD to abolish actin polymerisation. This revealed that AKMT mobility is tightly dependent on the integrity of PCKMT–FRM1–actin signalling, confirming that actin assembly is essential for AKMT redistribution.

### Apical methyltransferase forms a dynamic complex regulated by PCKMT

The residual redistribution of AKMT under CytD treatment suggested that a component of its apical dynamics may occur independently of full motility activation. To examine this dynamic behaviour directly, we performed fluorescence recovery after photobleaching (FRAP) experiments targeting the apical pools of AKMT and GAC. Both proteins recovered fluorescence after bleaching, indicating they are not stably anchored to the conoid but instead dynamically associate with the apical complex (Fig. 6d, e; Movie S7). Quantitative analysis showed that GAC-YFP recovered rapidly (with a half-time of fluorescence recovery (τ½) of 8.9 s and a maximum recovery level (Imax) of 78.1%) and AKMT-YFP recovered more slowly but to a similar extent (τ½ = 29.4 s, Imax = 73.4%). As expected for a stably integrated structural protein, IMC1-Halo showed negligible recovery. (Fig. 6f, g)

Attempts to perform FRAP on PCKMT-YFP and FRM1-YFP were unsuccessful due to the protein’s low expression levels: the YFP signal was barely detectable and photobleached before acquisition. Nevertheless, live imaging and fixed-cell analysis revealed no evidence of relocalisation following stimulation, suggesting that PCKMT is a stably anchored conoid component (Fig. 6a, (−Rapa); Supplemenatary Fig. 10f).

Together, these results indicate that apical proteins such as AKMT and GAC form a dynamic, reversible complex whose localisation depends on PCKMT-mediated actin polymerisation and underpins motility initiation.

## Discussion

Our findings demonstrate that two conoid-anchored, Apicomplexa-specific methyltransferases, PCKMT and AKMT, play distinct yet complementary roles in motility initiation. PCKMT is required for the recruitment and anchoring of FRM1, enabling actin filament formation and conoid extrusion, while AKMT^15^ ensures the recruitment of GAC, linking F-actin to surface adhesins for force transmission^9^. Together, these enzymes define a dual methylation system that coordinates cytoskeletal activation and mechanical coupling at the parasite apical complex, ensuring that motility is initiated only once both systems are properly engaged.

Strikingly, PCKMT operates with exceptional specificity: proximity labelling and methyl-enrichment identified only a limited number of candidate substrates, with FRM1 emerging as the sole effector critical for motility. Loss of PCKMT disrupts FRM1 recruitment to mature conoids, and mutation of its catalytic residue (H542A) mirrors this phenotype, blocking actin nucleation and gliding. These findings indicate that PCKMT’s enzymatic activity, and to a certain extent its structural presence, are both required to license motility initiation. This dual role as both scaffold and methyltransferase echoes that of large SET-domain proteins like KMT2D/MLL4, which stabilise interacting partners while methylating specific chromatin targets^29^. By analogy, PCKMT functions as a minimalist apical scaffold whose catalytic activity is channelled almost exclusively towards FRM1, creating a tightly focused methylation checkpoint at the onset of actin assembly.

In contrast to this narrow specificity, AKMT targets a broader set of substrates, as reflected by its wider distribution in microscopy and proteomic analyses. Consistent with previous work, AKMT relocalises from the conoid to the cytosol upon motility activation^15^, and our data show that this translocation requires both PCKMT and FRM1. Surprisingly, GAC was absent from our methylome enrichment assays, suggesting that it is not directly methylated but instead recruited via one or several methylated intermediates. Further work is needed to elucidate the mechanism of GAC recruitment to the apical complex. What is clear, however, is that both AKMT and GAC are recruited to the apical complex in immotile parasites and translocate towards the basal pole immediately after motility is initiated via actin dynamics. Whether actin itself, or an actin-associated protein, is among AKMT’s direct substrates linked to this translocation remains an open question.

In metazoan cells, several methyltransferases directly modify cytoskeletal proteins to regulate filament dynamics and cell motility. SETD3 methylates β-actin at His73, a modification that stabilises filaments and promotes motility^30,31^, while SETD2 and SET8 have been reported to methylate α- and β-tubulin as well as β-actin, influencing microtubule organisation and actin polymerisation^11,32,33^. Other enzymes, such as PRMT5 and EZH2, have also been implicated in actin methylation and cytoskeletal control^34–36^. In contrast, our data suggest that neither AKMT nor PCKMT directly methylates actin in *T. gondii*. Actin was not enriched in either our methyl-enrichment or TurboID assays, and no specific actin methylation signals were detected in the presence or absence of these enzymes. This implies that any impact of these apical methyltransferases on actin dynamics is likely indirect, possibly mediated through their primary effectors, such as FRM1 and other apical complex components, or that any direct actin methylation is transient or occurs at levels below detection in our current experiment setup. This strategy may reflect an adaptation to the parasite’s highly polarised cytoskeleton, allowing tight spatial and temporal control of actin assembly without altering core actin properties.

Interestingly, α- and β-tubulin appeared enriched in the wild-type but not in the AKMT-KO methyl-enrichment dataset. Tubulin methylation has been reported in other eukaryotes, and the lysine 40 (K40) residue of α-tubulin is a particularly well-conserved methylation site known to influence microtubule function in mammalian cells^33^. Consistent with this, previous proteomic analyses in *T. gondii* identified methylated peptides in both α- and β-tubulin isoforms^37^, supporting the possibility of direct methylation by AKMT. However, our recent mutagenesis of K40 in α-tubulin, designed to mimic (K40M) or prevent methylation (K40A), did not alter microtubule stability^38^. This suggests that the functional impact of α-tubulin methylation in *T. gondii* may not involve structural stabilisation but could instead fine-tune microtubule dynamics or interactions through more subtle regulatory mechanisms.

Together, these data support a model in which the core motility machinery is pre-assembled at the apical complex of non-motile parasites, poised for rapid activation upon stimulation (Fig. 7). Consistent with current models^1^, in resting parasites, PCKMT anchors FRM1 at the preconoidal rings, preparing the site for actin nucleation (Fig. 7; step 1). Upon stimulation, FRM1 promotes conoid extrusion and local actin polymerisation, facilitating engagement with GAC and the glideosome (Fig. 7; step 2). Concurrently, AKMT disengages from the conoid in an actin-dependent manner, likely methylating downstream targets that complete the transition to productive motility. FRAP analyses revealed that AKMT and GAC are not stably anchored but instead exchange dynamically between apical and cytosolic pools, maintaining the system in a “primed” state ready for immediate response to upstream signalling.

In the absence of PCKMT, FRM1 is not recruited to the conoid, preventing local actin nucleation. Nevertheless, AKMT and GAC are still independently recruited, but neither protein can translocate from the apical complex, resulting in a complete failure to initiate motility (Fig. 7; Step 3).

In *P. falciparum*, the PCKMT orthologue SET9 has so far been implicated only in gametocyte differentiation^22^. Our data now reveal that SET9 localises apically in segmented merozoites and that its depletion causes a strong invasion defect during the asexual blood stages, mirroring the phenotype observed in *T. gondii* PCKMT knockouts. Given that *Plasmodium* parasites possess a highly reduced conoid complex compared with other apicomplexans^39^, these findings raise intriguing questions about how SET9 is positioned within this simplified apical architecture and whether it contributes to motility or additional processes across the parasite life cycle, questions that are currently under investigation.

In summary, our data reveal a tightly ordered methylation–actin cascade that coordinates motility initiation (Fig. 7; step 4). PCKMT first licenses FRM1-mediated actin nucleation, which in turn enables AKMT translocation and GAC recruitment to transmit force. This stepwise sequence places lysine methylation at the core of motility control, providing a unifying framework for how apical signalling, cytoskeletal dynamics, and mechanical output are synchronised in *T. gondii* and likely conserved across apicomplexans.

## Materials and methods

### Culture conditions

Human foreskin fibroblasts (HFF) were cultured in Dulbecco’s modified Eagle’s medium (DMEM) supplemented with 10 % foetal bovine serum, 25 mg/mL gentamicin, and 2 mM L-glutamine until they reached 100 % confluency. Parasites were then allowed to infect the HFF monolayer and maintained at 37 °C with 5 % CO_2_ in an incubator.

### Generation of plasmids

Guide RNAs targeting the gene of interest were designed using EuPaGDT ^40^ and are listed in Supplemenatary Table S1. The sgRNAs were ligated into a vector coding for Cas9-YFP expression as previously described^41^. Briefly, gRNA sequences were ordered as single-stranded oligomers (ThermoFisher) with overhangs to allow for ligation to the Cas9-YFP vector^41^. Primers were diluted at equimolar concentrations in annealing buffer (10 mM Tris, pH 7.5–8.0, 50 mM NaCl, 1 mM EDTA), heated at 95℃ for 5 minutes and left to cool down for at least 2 hours. Cas9-YFP vector was linearised using BsaI (NEB). Annealed primers and digested vector were ligated using T4 ligase (NEB) overnight at room temperature. The ligated vector was transformed into DH5α bacteria, and colonies were screened for positive gRNA-containing vectors.

To generate the vectors containing the second copy of *pckmt* fragments containing promoter region, TPR domain, SET domain, Ankyrin domain, proline-rich motif, and 3’UTR region were amplified by PCR with overhangs to allow for insertion with Gibson assembly kit (NEB) as described in Supplemenatary Fig. 5. Insertion of the different fragments was verified by PCR.

### Generation of transgenic parasites

The CRISPR/Cas9 system was used to generate transgenic parasites. Repair templates for integrating a tag and a loxP sequence were generated following the method described by Stortz et al. ^2,17,42^. Briefly, for tagging, the repair templates were PCR-amplified from vectors carrying tags such as 3xHA, SYFP2, Halo, and SNAP. Primers used for this process were designed to bind the vectors and included 50 bp of homology to the gene. For the upstream loxP, oligonucleotides containing the loxP sequence, flanked by 33 bp of homology on both sides of the gene, were ordered as single-stranded DNA from Thermo Fisher.

The HaloTag system was employed to label PCKMT due to its versatility in conjugating different fluorophores and its ability to provide high signal-to-noise ratios and specificity when combined with Janelia dyes (Promega). This approach was particularly advantageous for imaging this low-expressing protein. Other proteins, such as FRM1 or AKMT, were tagged with YFP derivative (SYFP2), and GAC was tagged with mCherry or YFP.

Parasite transfection, sorting, and screening for positive clones were performed as described by Stortz et al. ^17^. In short, repair templates and Cas9-YFP expression vectors were co-transfected into freshly lysed or mechanically lysed RHDiCre Δ*ku80*^43^ tachyzoite parasites. After 24-48 hours post-transfection, parasites were sorted into 96-well plates using FACS (FACSARIA III, BD Biosciences). Single parasite clones were isolated and examined via genomic PCR to confirm correct modifications and sequenced where necessary.

When indicated in the figure legend, TLAP1 (TrxL1-associating proteins)^44^ endogenously tagged was used to show the parasite’s shape.

CBem and second copies of *pckmt* were integrated into the *uprt* (*uracil phosphoribosyltransferase*) locus by CRISPR/Cas9-mediated targeting. The corresponding amplicons, flanked by 50-nucleotide homology arms to the 5′ and 3′ untranslated regions (UTRs) of *uprt*, were inserted as previously described^2^. Parasites lacking UPRT activity were selected using 1µM 5-fluoro-2-deoxyuridine (FUDR; Thermofisher)^45^.

Primers are listed in Supplemenatary Table S2, and the transgenic parasite lines generated/used in this study are listed in Supplemenatary Table S3.

To induce KO, 50 nM rapamycin was added for one to two hours, followed by a wash with fresh DMEM media, and parasites were allowed to grow until fixation.

### Immunofluorescence assay (IFA) in *Toxoplasma*

HFFs were cultured on sterile coverslips in 24-well plates to establish a monolayer for parasite infection. The parasites were then allowed to infect the HFF monolayer for 24-48 hours. The coverslips were fixed with 4 % Paraformaldehyde (PFA) for 20 minutes, followed by block-permeabilisation using 3% BSA and 0.1% Triton-100 in PBS for 20 minutes. Primary and secondary antibodies of required dilutions were prepared in the block-permeabilising solution (See antibody list in Supplemenatary Table S4). The coverslips were mounted onto slides using ProLong™ Gold Antifade Mountant with DNA Stain DAPI (Thermo Scientific, USA) and imaged using a Leica DMi8 inverted live cell widefield microscope or an Abberior STED microscope.

To visualise Halo-tagged proteins, parasites were incubated with 4-12 nM HaloTag Janelia 646 (Promega, GA112A) overnight prior to fixation. For colocalization analysis, 50 nM HaloTag Janelia 646 was used. After dye application, parasites were washed three times with PBS and incubated in media for 10 minutes prior to fixation.

### Expansion microscopy

ExM was performed as described by Dos Santos et al.,^46^ with some modifications. Briefly, intracellular parasites on 12mm coverslips were fixed with 4% PFA, followed by three washes with PBS, and stored at 4°C. The fixed coverslips were then treated with a 1.4% formaldehyde and 2% acrylamide mix at 37°C for 5 hours, followed by a gelation step. A 35 µL drop of Monomer Solution (19% sodium acrylate, 10% acrylamide, 0.1% bis-acrylamide) supplemented with 0.5% Tetramethylethylenediamine (TEMED) and 0.5% Ammonium persulfate (APS) per coverslip was used for gelation on ice in a humid chamber for 5 minutes, then at 37°C for 1 hour. After gelation, the gel was denatured with denaturation buffer (200 mM SDS, 200 mM NaCl, 50 mM Tris in ultrapure water, pH 9) at 80/85°C for 1.5 hours. Gels were then expanded overnight in deionized water (dH_2_O) at 4°C. A small piece of the gel was cut and subjected to immunostaining. For immunostaining, antibodies were diluted in freshly prepared PBS with 2% BSA. The gel was incubated with primary antibodies for 3 hours at 37°C, followed by four washes with PBS containing 0.2% Triton X-100 (PBS-Tx100). The gel was then incubated with secondary antibodies for 2.5 hours at 37°C or overnight at 4°C, followed by an additional 2 hours at 37°C, and then washed four times with PBS-Tx100. Antibody concentrations are listed in Supplemenatary Table S4. The gel was fully expanded in dH_2_O for a few hours or overnight. To stain the nuclei, 0.8 µM Hoechst 33342 (diluted in dH_2_O) was then used for 3 minutes, followed by one wash. After determining the orientation of the gel, it was mounted on a glass-bottom dish pre-coated with poly-L-lysine. Widefield images were acquired in Z-stacks with 0.3 µm increments using a Leica DMi8 wide-field microscope.

### Sample preparation for cryo-ET

Human foreskin fibroblasts infected with *T. gondii* from the loxP*pckmt*-Halo strain were treated with 50 nM rapamycin for 72 h to induce the knockout. The uninduced (wild-type) control sample originated from an unpublished dataset prepared for a previously published manuscript^3^, using the loxP*cgp*-Halo/*pcr4*-Halo strain that was, apart from the induction, prepared in the same way as iKO. On the day of harvesting, parasite-infected fibroblasts were stained with 0.04 µM JaneliaFluor 646 Halo dye (Promega, GA1120) for 1h at 37°C. Cells with parasites were scratched from the dishes, centrifuged for 5 min at 800 g and resuspended in fresh DMEM media. Samples were then syringed three times and centrifuged for 5 min at 600 g. Parasites were resuspended in 0.5 mL PBS and stained with NucBlue (Hoechst 33342, Thermo Fisher Scientific) for 15 min at 37°C. Parasites were washed with PBS and treated with +/- 2 µM calcium ionophore (Sigma-Aldrich A23187) for 5 min at RT and washed with PBS. Carbon lacey cryo-EM grids with mesh 300 (Quantifoil) were plasma-cleaned for 30s with a 90:10 argon: oxygen mixture using plasma cleaner (Fischione 1070) set to 100% power and flow of 30 SCCM. Subsequently, 4 µL of parasites were applied on the grid mounted on GP1 plunge freezer (Leica), backside blotted for 2-3 s and plunge frozen into liquid ethane. The environment in the plunger was set to 99% humidity and 22°C. Vitrified grids were clipped into Autogrids in liquid nitrogen vapours and stored in liquid nitrogen.

### Cryo-electron tomography

Grids with vitrified parasites were loaded onto a Titan Krios G4 electron microscope (Thermo Fisher Scientific) operated at 300 kV, equipped with Falcon4i camera and Selectris X energy filter. Low magnification images were taken to assess ice thickness and parasite distribution over different grid squares. Then, medium magnification montages of the grid squares were acquired to identify apical sites of the parasites. Tilt series were acquired using SerialEM^47^ with a dose symmetric tilting scheme^48^, range of +/-60° with 2° increment, at pixel size of 3.033 Å, exposure time of around 1 s with exposure dose of ∼2.3 e⁻/Å^2^ that accumulated to around 140 e⁻/Å^2^ throughout the entire tilt series.

### Tomogram reconstruction

Frames acquired in the EER format were converted to TIFF, followed by motion correction, CTF estimation and tilt selection using Relion-5^49^. Tilt stacks were aligned using the AreTomo2 wrapper implemented in Relion-5. Tomograms were reconstructed with stand-alone AreTomo2^50^. Reconstructed tomograms were then subjected to CTF-deconvolution from IsoNet^51^ to improve visualization.

### Segmentation and visualization

Reconstructed tomograms were segmented using MemBrain-seg^52^ and Dragonfly (Comet Technologies Canada Inc. Dragonfly 3D World (v. 2024.1), Comet Technologies Canada Inc., https://dragonfly.comet.tech/). Firstly, membrane segmentation was predicted using MemBrain-seg (using model MemBrain_seg_v10_beta.ckpt) and imported to Dragonfly. Then, non-membranous structures were segmented manually using either full or Otsu thresholding during segmentation. Classes of segmented organelles were exported as TIFF files and a 3D Gaussian filter from FIJI^53^ was applied before importing to ChimeraX^54^ for visualization together with the reconstructed tomograms. Movies of tomograms and segmentations were generated using custom Python scripts.

### Plaque assay

Parasites were released by passing them through a 26-G needle. 1000 parasites per condition were allowed to infect the HFF monolayer cultured on a 6-well plate. The parasites were left undisturbed at 37°C and 5% CO_2_ for 6-7 days to form plaques. The cells are then washed with PBS and fixed with ice-cold 100 % Methanol (-20 °C) for 20 minutes and later stained with Giemsa stain to visualise the plaques. The plaques were imaged using a Leica DMi8 (Leica, Germany) inverted live cell widefield microscope.

### Invasion assay

Parasites were pre-induced ± 50 nM Rapa for 72 h. 5 × 10⁶ freshly egressed tachyzoites were resuspended in 2 mL DMEM and kept on ice for 10 min before inoculation. Parasites were then allowed to invade confluent HFF monolayers for 1 h at 37 °C, then fixed with 4% PFA for 10 min and washed three times with PBS. To distinguish extracellular and intracellular parasites, sequential immunofluorescence staining was performed. For extracellular labelling, samples were blocked without permeabilisation and incubated with anti-SAG1 for 1 h, followed by secondary antibody. After PBS washes, intracellular parasites were visualised by blocking buffer with 2% BSA, followed by staining with anti-IMC1(plus secondary) to label all parasites. Samples were imaged by a Leica DMi8 widefield microscope. A minimum of 100 parasites were scored per condition to calculate invasion efficiency, expressed as the percentage of intracellular (dual-labelled) versus extracellular (single-labelled) parasites.

### Replication assay

Parasites were allowed to infect HFF monolayer-seeded glass coverslips for one hour. The non-invaded parasites were then washed off with PBS to synchronise the replication of parasites within the host cells. DMEM media (Sigma) was added to the coverslips, and the parasites were incubated at 37 ℃ for 24 hours to allow for replication within the host cells. Induced KO for loxP-*pckmt-halo* were pre-incubated with 50nM rapamycin for 72 hours prior to the infection of coverslips.

The coverslips were then fixed, and the standard IFA protocol was followed as described previously using anti-GAP45 (1:5000), to visualise the parasites as the primary antibody. The parasites were imaged using the Leica DMi8 inverted live cell widefield microscope (Leica, Germany), and the number of vacuoles in the various cell stages of the parasites was determined for each mutant strain. To improve the statistical significance of the observations, three biological replicates were performed.

### Live Gliding assay

Pre-warmed FBS was added onto the surface of a Cellvis 29 mm glass-bottom dish (Cellvis, USA) and incubated for 1-2 hours and then aspirated to provide a surface to facilitate parasite gliding. Parasites were harvested and passed through a 3µm filter to remove debris. The parasites were pelleted at 1500 g for 5 minutes and washed with a pre-warmed gliding buffer to remove media. They were then resuspended in the gliding buffer (1mM EGTA and 100mM HEPES in HBSS solution), ensuring a concentration of 4×10^6^ parasites/mL. The parasites were added to the FBS-coated glass-bottom dishes. Gliding efficiency was recorded using the Leica DMi8 inverted live cell widefield microscope with ×100 oil objective by capturing the gliding of the parasite every second for a total of 10 minutes in the DIC channel. In the case of iKO parasites, a stack image was performed after each video to determine the efficiency of our protein of interest (POI) depletion. Only parasites lacking the analysed POI were considered for the quantification of gliding motility for iKO parasites; only parasites with movement were measured for speed and distance.

The distance travelled by the parasites (≥9 parasites per replicate) as well as the speed at which the gliding motion occurred were assessed using the ManualTracking Plugin in Image J across three independent biological replicates.

### Egress assays

To investigate PCKMT behaviour during egress, loxP*pckmt*-Halo/CbEm parasites induced with ± 50 nM Rapa for 54 h before triggering egress, transiently transfected with the vector pTub-pTub-sag1ΔGPI-dsRed and grown in HFFs on a glass-bottom dish for approximately 30 hours before induction of egress with 4 µM Calcium ionophore A23187(CI) (Sigma-Aldrich, C7522-1mg). Images were taken in auto-focus mode with a 100x oil objective lens. All videos were recorded in triplicate per condition at a minimum. Live microscopy was performed using a Leica DMi8 widefield microscope equipped with a DF C9000 GTC camera in a heated chamber with 5% CO_2_.

For the fixed egress assay, parasites were pre-induced ± 50 nM Rapa for 54 h, and Halo dye was added to live cultures 2-4 h before fixation. Egress was triggered with 4 µM CI for 5min, and then fixed with PFA. Without permeabilisation, samples were blocked and stained with anti-SAG1 to label the extracellular parasites, followed by the corresponding secondary antibody. After permeabilisation with Triton X-100, an IMC antibody (plus secondary) was applied to label all parasites; an auto 488 antibody was used to anti-SYFP2. To block F-actin polymerisation, 2 µM Cytochalasin D(CytD) was added for 1h prior to egress induction. Antibody details are provided in Supplemenatary Table S4.

### Protrusion assay

LoxP*pckmt*-Halo, loxP*pckmt*-Halo/pckmt^com^-SYFP2, loxP*pckmt*-Halo/*pckmt*^H452A^-SYFP2 and loxP*frm1*-Halo. Parasites were pretreated ± 50 nM rapamycin for 72h prior to the assay. Parasites were mechanically released, filtered, pelleted, and resuspended in pre-warmed DMEM (without FBS) containing 4 µM CI. They were then added to 24-well plates containing poly-L-lysine-coated coverslips and subjected to brief centrifugation at 1000 rpm for 1 minute. After incubation at 37°C for 8-10 minutes to promote extrusion, parasites were fixed with PFA. The parasites were then processed for ExM and stained with anti-acetylated tubulin and Halo dye. Three biological replicates were performed, with at least 100 parasites counted per condition and three biological replicate were performed. The Halo antibody did not work effectively to detect the direct absence of PCKMT and FRM1 from the conoid.

### Fluorescence Recovery After Photobleaching (FRAP)

Intracellular *Toxoplasma gondii* parasites expressing YFP-tagged proteins (GAC-YFP, AKMT-YFP, MyoH-YFP) were seeded in µ-dishes (Ibidi) and cultured for 24 hours prior to imaging. Before FRAP, the medium was replaced with FluoroBrite™ DMEM (ThermoFisher) supplemented with 10% fetal bovine serum (FBS), L-glutamate, and gentamycin. Imaging was performed using a Leica DMi8 widefield microscope equipped with a FRAP module and a 488 nm laser.

Fluorescence was recorded every second for 10 seconds before photobleaching. Bleaching was performed at the apical tip of single parasites using a high-intensity pulse of the 488 nm laser, and fluorescence recovery was monitored every second for 60 seconds. Experiments were performed in biological triplicates, and at least six vacuoles were analysed in total across the replicates.

Image analysis was conducted using Fiji (ImageJ). Intensities were measured in the bleached area, a non-bleached parasite in the same field of view, and the background. Signals were corrected by subtracting background fluorescence and normalised using the non-bleached parasite to account for overall photobleaching. Recovery curves were generated from the normalised values. Data were then transformed so that the pre-bleach average intensity corresponded to 100%, ensuring comparability across samples with slightly different bleaching depths. To determine the characteristic recovery time (Tau, **τ**), the normalized post-bleach data were fitted to a single exponential function of the form:

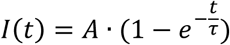

Where I(t) is the normalized intensity at time t, *A* is the plateau (maximum recovery) and τ is the time constant representing the time required to reach ∼63% of the final recovery level. From this fit, the half-time of recovery (t₁/₂) was also calculated using the relationship:

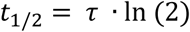

which corresponds to the time at which fluorescence intensity reaches 50% of the recovery plateau.

### Enrichment of methylated proteins and pull-down

Roughly 10^8^ parasites of the strains loxP*pckmt*-*Halo*, loxP*akmt*-*syfp2* and double AKMT/PCKMT inducible KO ± rapamycin for 72h were collected per technical (n= 2) and biological replicates (n =3; total samples = 6 per strain/condition). Parasites were released mechanically and filtered through a 3 µm membrane (Millipore). Parasites were washed with ice-cold PBS. Pelleted parasites were resuspended in 1 ml RIPA buffer (0.5% sodium deoxycholate, 150mM NaCl, 1mM EDTA, 0.1% SDS, 50mM Tris-HCl (pH 8.0), 1% Triton TX-100) with Pierce™ protease inhibitors and incubated at 4 °C for 1h.

During this time, 100 µl of Protein G Dynabeads™ (Thermofisher) per sample were each conjugated with 9.4 µg of αH4K20m3 antibody (Abcam) diluted in 200 µL PBS with 0.02% Tween™20 (Sigma) for 10 minutes at room temperature, followed by 3 washes in PBS with 0.02% Tween™ and a final wash in RIPA buffer.

Lysed parasites were incubated for 1 h with the conjugated magnetic beads. After that, the beads were then washed five times with 1 mL of RIPA buffer without Triton TX-100 but with protease inhibitor, followed by three washes with 50 mM Tris-HCl (pH 8). Washing steps were performed on ice. 10 µl samples of lysed parasites prior to incubation with beads, output after incubation and last wash together with 10% of the washed beads were preserved for Western blot analysis; the rest of the beads were pelleted and stored at −80 °C before being sent for mass spectrometry.

### Proximity labelling assay

10^8^ parasites of the strains *pckmt*-*TurboID, akmt*-*TurboID* and RHΔ*Ku80*DiCre (wild-type parasite) pre-treated for 2 or 6 hours with or without 150 μM biotin were collected following the protocol described above in biological replicates (n=4). Additionally, 10^7^ parasites treated with ± 150 μM biotin were collected for biotinylation analysis via Western blot. Pelleted parasites were also lysed with RIPA buffer with protease inhibitors for 1h. The supernatant from the lysis was incubated with 100 μL of beads per sample (Dynabeads™ MyOne™ Streptavidin T1, Invitrogen), prewashed with PBS, for 30 minutes at room temperature while being gently rotated. The beads were then washed five times with 1 mL of RIPA buffer without Triton TX-100 but with protease inhibitor, followed by three washes with 50 mM Tris-HCl (pH 8). Washing steps were performed on ice. While 10% of the beads were preserved for Western blot analysis, the rest of the beads were pelleted and stored at −80 °C before being sent for mass spectrometry.

### Protein quantification by LC-MS

Immunoprecipitated proteins were quantified by LC-MS essentially as described previously, with minor modifications^55^. Briefly, beads were incubated with 10 ng/µL trypsin in 1 M urea and 50 mM ammonium bicarbonate (NH₄HCO₃) for 30 minutes, followed by a wash with 50 mM NH_4_HCO₃. The supernatant was then digested overnight in the presence of 1 mM DTT. After digestion, peptides were alkylated and desalted prior to LC-MS analysis. For LC-MS/MS, peptides were loaded onto an Ultimate 3000 RSLCnano system and separated on a 25 cm analytical column (75 µm inner diameter, 1.6 µm C18, Aurora-IonOpticks) using a 50-minute gradient from 2% to 35% acetonitrile in 0.1% formic acid.

The effluent from the HPLC was directly electrosprayed into an Orbitrap Exploris 480 (Thermo) operated in data-dependent mode to automatically switch between full scan MS and MS/MS acquisition. Survey full scan MS spectra (from m/z 350-1200) were acquired with resolution R=60,000 at m/z 400 (AGC target of 3×106). The 20 most intense peptide ions with charge states between 2 and 6 were sequentially isolated to a target value of 1×105, and fragmented at 30% normalised collision energy. Typical mass spectrometric conditions were: spray voltage, 1.5 kV; no sheath and auxiliary gas flow; heated capillary temperature, 275°C; intensity selection threshold, 3×10^5^.

Protein identification and quantification were performed with MaxQuant v2.4.1.0 using iBAQ-based quantification. Searches were carried out against the Toxoplasma gondii protein database (Uniprot_UP00000152_Toxoplasmagondii_Me49_20221024-08.04.16.64.fasta) with the following parameters: precursor mass tolerance 10 ppm, fragment mass tolerance 20 ppm, peptide FDR 0.1, protein FDR 0.01, minimum peptide length 7, fixed modification: carbamidomethylation (C), variable modification: oxidation (M). Razor and unique peptides were used for quantification, with a minimum of one peptide per protein and at least two ratio counts.

Quantified protein values (iBAQ, Z-score normalised) were compared using the adjusted t-test implemented in Perseus (v.2.1.0)^56^. Missing values were imputed from a normal distribution (width = 0.3, downshift = 4). Differential enrichment analysis was performed using a permutation-based false discovery rate (FDR) approach, with thresholds set at 0.05 for AKMT and DAMT-iKO, and 0.1 for PCKMT in the methylation enrichment assay. For the proximity labelling experiment, an FDR threshold of 0.08 was applied for PCKMT. In all cases, a constant *s₀*parameter of 0.1 was used to stabilise variance estimation.

Volcano plots depicting the different abundance of proteins detected were constructed using R (v. 4.5.1) and RStudio (2025.09.1 Build 401).

### *Plasmodium falciparum* Parasite Culture and Transfection

Asexual parasites were maintained in RPMI 1640 with AlbuMAX (Gibco, Thermo Fisher Scientific) and supplemented with 2.5mM L-glutamine and 25µg/mL Gentamicin. Parasite cultures were incubated at 37°C in a 94% N_2_, 5% CO_2_, and 1% O_2_ gas mixture. Blood smears were routinely made and stained with Hemacolor® Rapid staining of blood smear (Sigma-Aldrich) to determine the parasitemia of the *Plasmodium* cultures. 10µg of plasmid DNA was transfected into mature schizonts of DiCre-expressing 3D7 parasite line (B11DiCre)^57^, isolated using 60% Percoll (Sigma-Aldrich), using the Amaxa P3 Primary cell 4D-Nucleofector® X Kit L (Lonza). 24 hours post-transfection, parasites maintaining the plasmid episomally were selected using 2.5nM WR99210 (Jacobus Pharmaceutical Company). The parasites were then subjected to 400µg/ml G418 (Sigma-Aldrich) to select for genomic integration of the construct. The media and drugs were replenished every 2 days. Genomic DNA was isolated from 200µL of infected red blood cells (iRBCs) using the Qiagen Blood and Tissue kit. An insertion PCR was conducted to verify the genomic integration using the appropriate primers (Supplemenatary Table S2).

Synchronous cultures were obtained by isolating mature schizonts using 60% Percoll (Sigma-Aldrich), following which they were allowed to infect fresh RBCs for 5 hours. Subsequently, 0-5-hour-old rings were obtained using 5% D-Sorbitol solution (Sigma-Aldrich). To understand the role of PfSET9 in the parasites, KO was induced at the various asexual blood stages of the synchronised parasite culture- at rings (0 hours), trophozoites (24 hours), and schizonts (42 hours), using 100nM Rapamycin (Sigma-Aldrich) while maintaining a non-induced, synchronised parasite population as a control.

### Immunofluorescence assay (IFA) in *Plasmodium*

Immunofluorescence analysis was performed as described in previous studies^58,59^. Thin blood smears were created on microscopic slides and fixed with 4% paraformaldehyde (PFA) in PBS for 10 minutes. The fixed smears were washed three times with 1x PBS. The parasites were permeabilised with 0.1% Triton X-100 in PBS for 10 minutes, followed by 3 washes with 1x PBS. The parasites were blocked overnight with 3% BSA in PBS at 4℃. The parasites were incubated with anti-HA (1:500, Rat, Roche) diluted in blocking solution for 1 hour at room temperature. The unbound primary antibody was removed with three 5-minute washes with PBS. The parasites were then incubated with appropriate secondary antibodies (Supplemenatary Table 5) diluted in blocking solution for 45 minutes at room temperature. The washing steps were repeated as described previously. All the steps described were performed in a humid chamber. A couple of drops of ProLong™ Gold Antifade mountant with DAPI (Invitrogen) were added before covering the slides with 24 x 60mm coverslips.

## Data analysis and statistics

Two-tailed Student t-test and one-way ANOVA were used to determine the statistical significance between groups.

## Software and online applications

All sequences were obtained from the ToxoDB database^60,61^. Protein structure predictions and models were generated using ColabFold v1.5.2^62^, Alphafold3^63^, and ChimeraX (v. 1.9 ^64^). sgRNA designing was aided by the Eukaryotic Pathogen sgRNA Design Tool (EuPaGDT) ^65^. *In silico* cloning was conducted using ApE (v. 3.1.3 ^66^) and Benchling (https://benchling.com). BioEdit (v.7.0.5.3 ^67^) was used in the analysis of sequencing results. Image acquisition was carried out using Leica LAS X software (v. 3.10.0.28982; RRID: SCR_013673) for the Leica DMi8 inverted live cell widefield microscope and Imspector (v.16.3.19714-w2408) for the Abberior STED microscope. All image processing and analysis were done using ImageJ (v. 2.16.0/1.5p)^53,68^. Cryo-ET data were acquired using SerialEM (v.4.1.1)^47^. Relion-5 (v. 5.0.0)^49^ and AreTomo2 (v. 1.1.2)^50^ were used for tomogram reconstruction and IsoNet (v. 0.2.1)^51^ for CTF-deconvolution. Tomogram segmentation was performed using MemBrain-seg (v. 0.0.5)^52^ and Dragonfly (Comet Technologies Canada Inc. Dragonfly 3D World (v. 2024.1), Comet Technologies Canada Inc., https://dragonfly.comet.tech/). Visualisation of tomograms together with segmentations was prepared using FIJI/ImageJ (v. 2.16.0/1.54p)^53^, ChimeraX (v.1.8)^54^ and custom Python (v. 3.13.1) scripts. Inkscape v1.4 was employed to generate the figures for this manuscript (www.inkscape.org). RStudio (v. 4.4.1 and 4.5.1) was utilised to perform statistical analysis and data visualisations.

## Authors contributions

P.Q. performed the characterisation of PCKMT, generation of mutants and analysed the data. T.K.: performed all experiments on *P. falciparum*. O.K. performed cryo-ET, analysed the cryo-ET data and wrote the cryo-ET results section. W.L. created some of the strains used in this publication and performed some imaging assays. I.F. performed the LC-MS and conducted the relevant analysis. S.M. analysed the cryo-ET data and contributed resources. E.J-R. designed and coordinated the project and experiments, analysed the data, wrote the paper and contributed with resources.

## Acknowledgements

We thank all colleagues who generously provided antibodies and reagents for this study. We are particularly grateful to Prof. Markus Meissner for his mentoring and his lab for all the insightful discussions that helped improve this work, and to Dr. Tobias Spielmann for his guidance in establishing *Plasmodium falciparum* culture in our laboratory and for training T. K. during the initial setup. P.Q. acknowledges support from the CSC Fellowship (grant number 202106300007). This work was supported by the DFG Priority Programme SPP2225 “EXIT Strategies of Intracellular Pathogens” (Project JI 463/2-2) and by a DFG Equipment Grant (INST 86/1831-1).

## Data availability

All parasite strains and imaging data generated for this paper are available from the authors on reasonable request.

## Movie captions

**Movie S1:** Cryo-ET and segmentation of wild-type parasites.

**Movie S2:** Cryo-ET and segmentation of PCKMT-depleted parasites.

**Movie S3:** Reconstructed tomograms from Supplemenatary Fig.S2 b-e

**Movie S4:** Gliding movies of loxP*pckmt*-Halo parasites with or without induction 72h prior to recording.

**Movie S5:** Egress movie showing loxP*pckmt*-Halo expressing SAG1ΔGPI-dsRed (in magenta) to mark PV space and Chromobody-emerald (Cbem; in yellow) to show intravacuolar network.

**Movie S6:** Gliding motility in the complemented and catalytically inactive second copy of PCKMT.

**Movie S7:** FRAP analysis of AKMT-YFP, GAC-YFP and IMC1-Halo.

