## Supplementary Figures for "Dual apical methyltransferases orchestrate motility initiation in apicomplexan parasites"

[1] Experimental parasitology. Faculty of Veterinary Medicine. Ludwig-Maximillian-University (LMU). Munich Germany

[2] European Molecular Biology Laboratory (EMBL), Molecular Systems Biology Unit, Heidelberg, Germany

[3] Collaboration for joint PhD degree between EMBL and Heidelberg University, Faculty of Biosciences

[4] Department of Parasitology, College of Veterinary Medicine, Sichuan Agricultural University, Chengdu, China

[5] Protein Analysis Unit, Faculty of Medicine, Biomedical Center (BMC), Ludwig-Maximilians-University (LMU), Munich, Germany

[6] European Molecular Biology Laboratory, EMBL Imaging Centre, Heidelberg, Germany

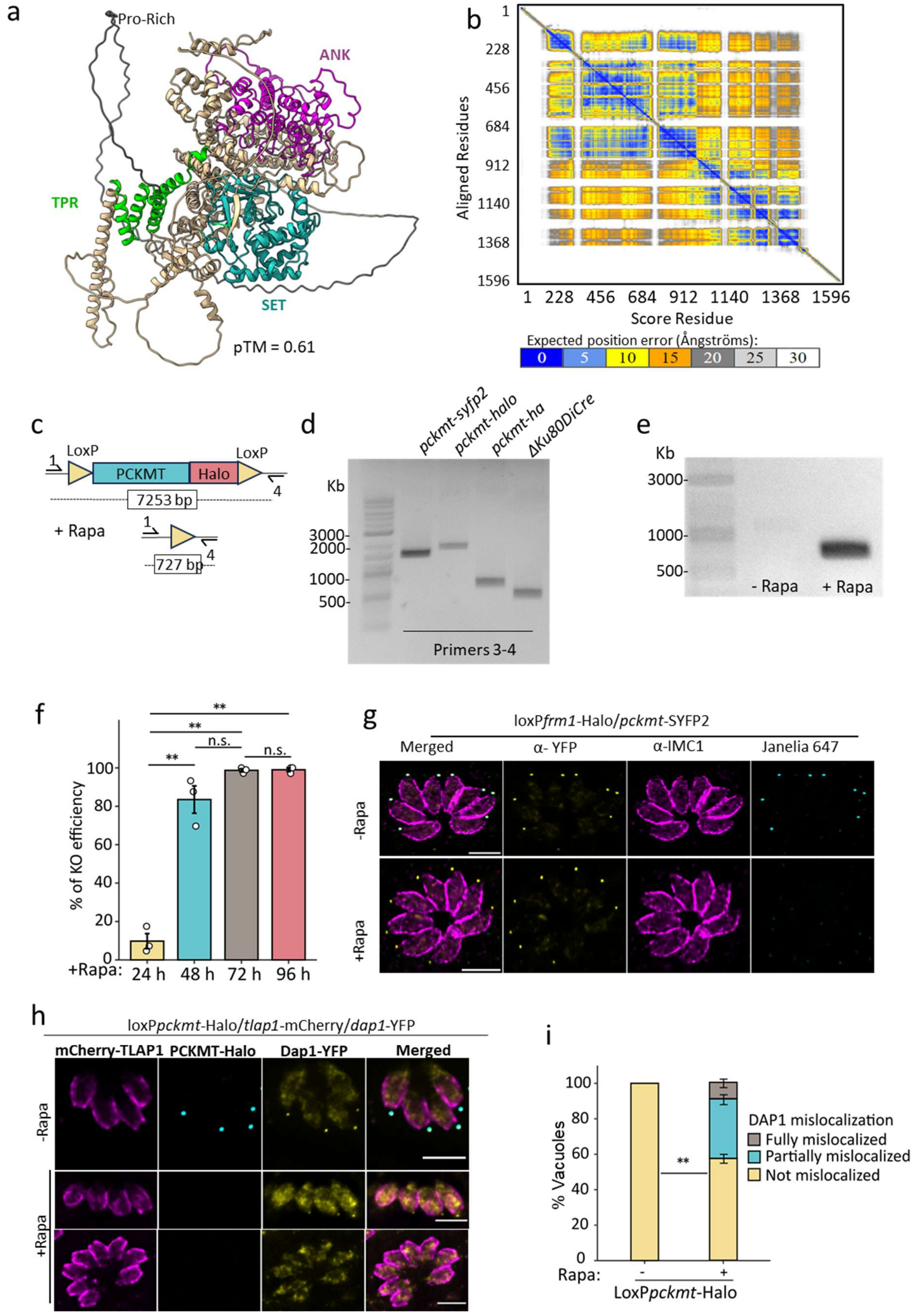


**Supplementary Fig. 1 | Structure prediction and genetic analysis implicate PCKMT in conoid organisation and selective anchoring of FRM1 and DAP1.**

***(a)*** *AlphaFold2*-predicted structure of PCKMT showing the ordered domains, TPR repeats (green), SET catalytic core (cyan), and ankyrin repeats (magenta), connected by extended linkers predicted with low confidence (beige). The predicted template modelling score (pTM = 0.61) indicates that while the overall domain topology is likely correct, the relative domain orientation may be uncertain.

**(b)** Predicted aligned error (PAE) heat map from *AlphaFold3* (accessed via https://alphafoldserver.com) for full-length PCKMT. The model reveals four well-defined regions: TPR, ankyrin, SET, and a proline-rich C-terminal segment. Colours indicate expected positional error between residue pairs (0–30 Å; blue = high confidence, yellow = low). The continuous low-PAE diagonal and distinct blue blocks correspond to intra-domain regions of high structural confidence, whereas warmer inter-domain regions denote flexible or poorly defined relative orientations.

**(c–e)** Schematic representation of the engineered *pckmt* locus (c), analytical PCRs confirming successful insertion of SYFP2, Halo, and HA tags (d), and excision of *pckmt* following 50 nM rapamycin treatment (e).
**(f)** Quantification of knockout (KO) efficiency at different time points after induction with rapamycin. Bars represent mean ± standard deviation (SD), with dots indicating the mean of each biological replicate (n = 3).

**(g)** Immunofluorescence analysis of PCKMT localisation in *loxPfrm1* parasites shows that PCKMT localises to the conoid independently of FRM1.

**(h, i)** Immunofluorescence analysis of DAP1 localisation in *loxPpckmt* parasites shows partial DAP1 mislocalisation upon PCKMT depletion (h). (i) Quantification of DAP1 mislocalisation after 72 h of rapamycin treatment (50 nM). Data represent mean ± SD from three biological replicates.

Statistical significance in all quantifications was assessed by a two-tailed Student's *t*-test (n.s., not significant; $^{**}p < 0.01$).

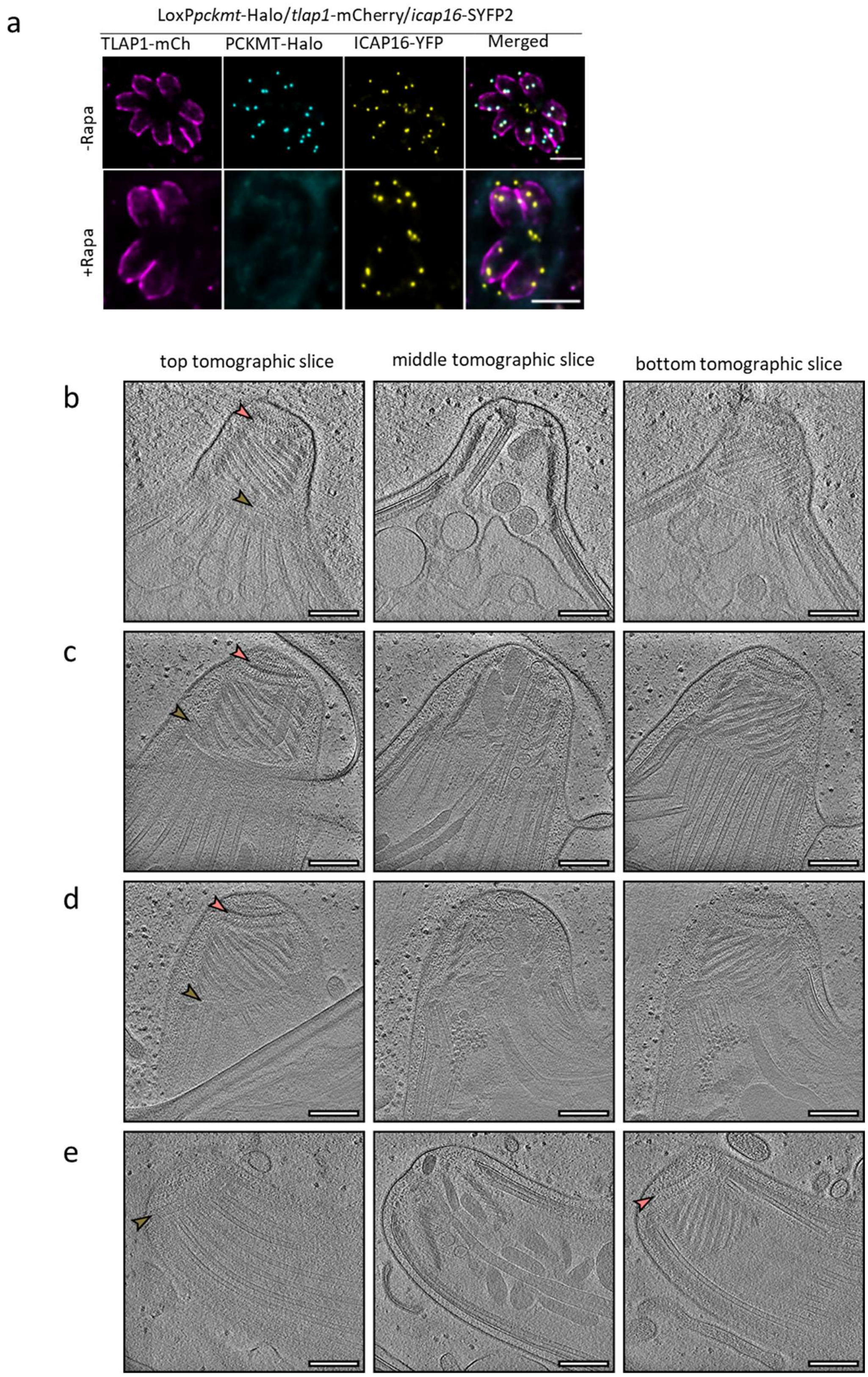


**Supplementary Fig. 2 | ICAP16 localises independently of PCKMT, and cryo-ET shows preserved preconoidal ring/APR architecture in extruded *T. gondii* PCKMT iKO parasites**.

**(a)** Immunofluorescence analysis of ICAP16 localisation in loxP*pckmt* parasites shows ICAP16 is independent of PCKMT.

**(b-e)** Reconstructed tomograms of *T.gondii* parasites presented as three example slices coming from the top, middle, and bottom of the reconstructed volume. Pink arrows indicate the PCRs, while khaki arrows indicate the APR. Scale bar: 200 nm. For full tomographic reconstructions, see Supplementary Video 3.

**(b)** Tomogram of extruded WT parasite (loxP*cgp*-Halo/*pcr4*-Halo strain). **(c)** and **(d)** Tomograms of extruded iKO parasites (loxP*pckmt*-Halo strain). **(e)** Tomogram of non-extruded PCKMT iKO parasites (loxP*pckmt*-Halo strain).

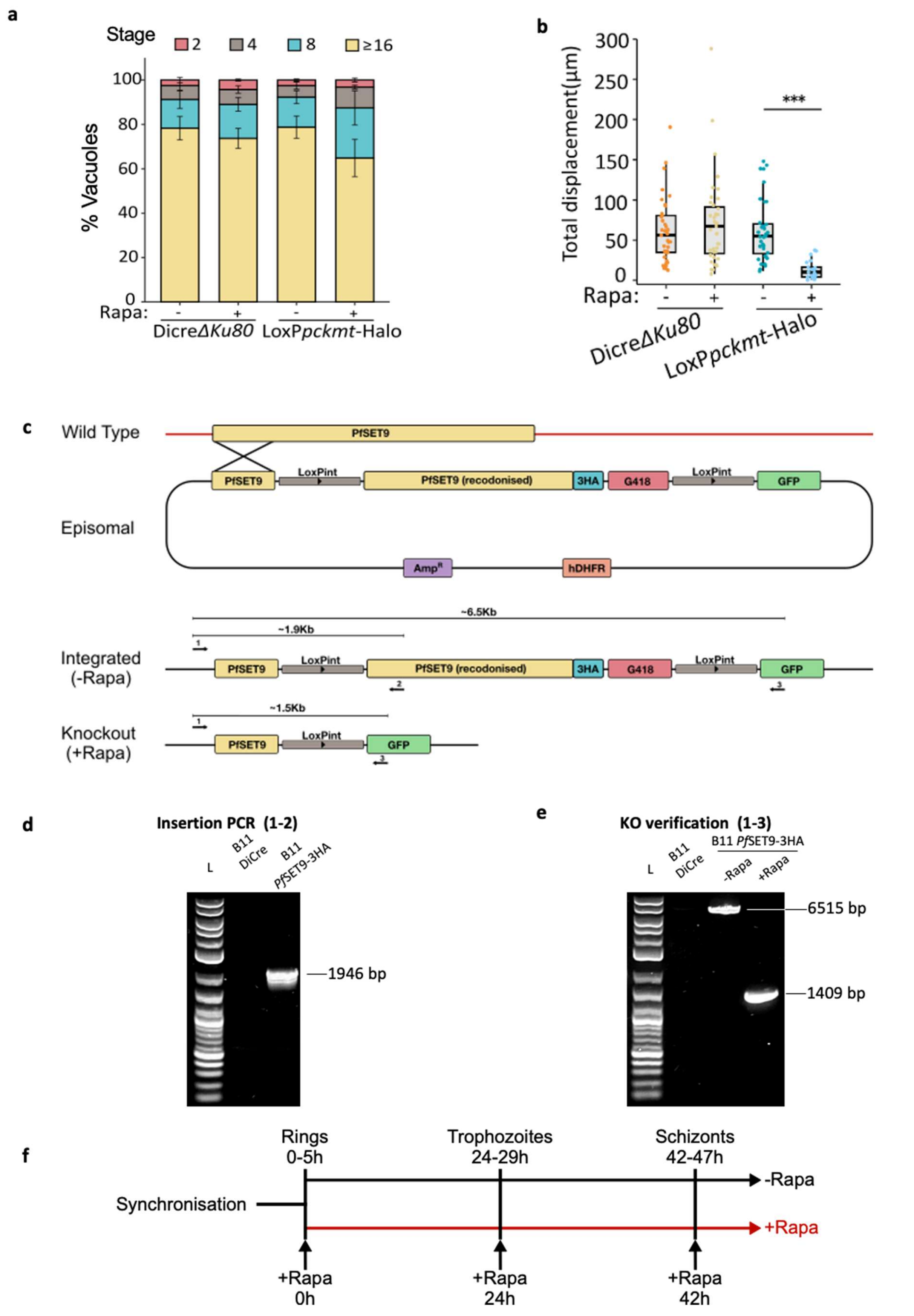


**Supplementary Fig. 3 | Functional assays and genetic engineering: stable replication with PCKMT perturbation, compromised gliding motility, and construction/verification of PfSET9 inducible knockout.**

**(a)** Quantification of the replication assay shows that PCKMT has no effect on replication. *T. gondii* parasites were pre-induced ± 50nM Rapa for 72 h and grown on HFF monolayers for 24 h. Vacuoles containing 2, 4, 8, or 16 parasites were counted; ≥100 vacuoles were counted in each replicate (n = 3).
**(b)** Gliding-distance analysis. Gliding tracks were extracted from time-lapse videos, and distances were measured in ImageJ. Parasites were pre-induced ± 50nM Rapa for 72 h. For each condition, ≥9 parasites per replicate were analysed across three independent biological replicates. Dots represent individual parasites; bars show the median. Statistical significance was assessed with an unpaired, two-tailed Student's t-test; ***$p < 0.001$.
**(c-f)** Schematic overview of the generation and validation of PfSET9-3HA inducible knockout (iKO) in *Plasmodium falciparum*.
**(c)** Schematic representation of the generation of a PfSET9-3HA iKO in *Plasmodium falciparum*. The scheme depicts the wild-type locus of PfSET9 along with the episomal plasmid containing the intended modifications to the wild-type locus to obtain a rapamycin-inducible transgenic line. The loci resulting from successful genomic integration and rapamycin-induced excision are also shown. The primers 1, 2, and 3 were used for insertion and knockout (KO) verification.
**(d)** Verification of integration of the construct into the wild-type locus using primers 1 and 2. B11 DiCre: 3D7 parental line expressing the DiCre system
**(e)** Verification of knockout generation 48 hours post-induction with 100 nM rapamycin.
**(f)** Schematic illustrating the rapamycin-mediated KO induction of tightly synchronised (0-5 hours) parasites at the various intraerythrocytic stages of the parasite (+ Rapa, red line). Non-induced, synchronised parasites (-Rapa, black line) were used as a control. Black arrows indicate the time points of induction with 100 nM rapamycin.

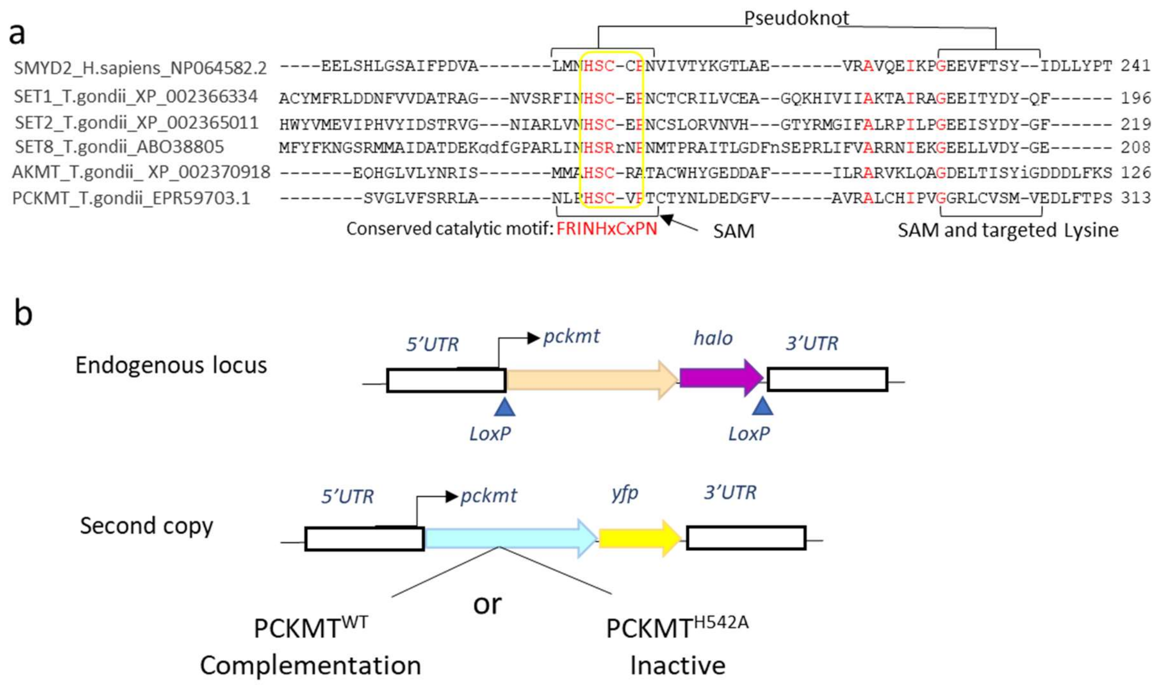


**Supplementary Fig. 4 | Conservation of the PCKMT catalytic SET domain and generation of a complementation system for functional analysis.**

**(a)** Multiple sequence alignment of the SET domain from *T. gondii* methyltransferases showing conservation of the catalytic motif HxC in PCKMT.

**(b)** Schematic of the complementation construct in which *pckmt–syfp2* was integrated into the *UPRT* locus under the control of the endogenous *PCKMT* promoter in the inducible *loxPpckmt–Halo* strain.

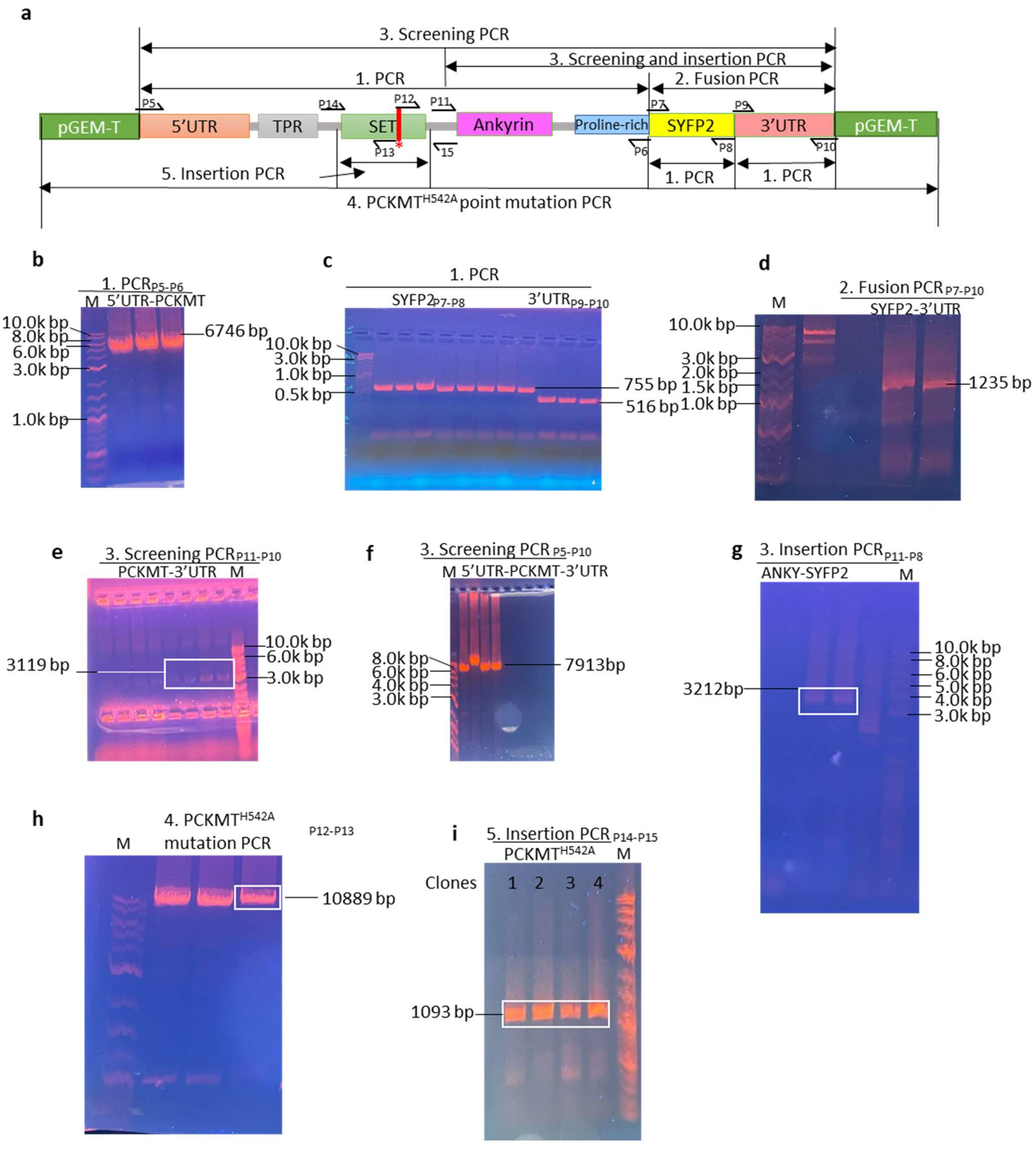


**Supplementary Fig. 5 | Generation and verification of PCKMT complementation and mutant constructs.**

**(a–f)** Construction and validation of the *pGEMT-PCKMT-SYFP2* complementation vector. **(a)** Schematic of cloning strategy showing insertion of *PCKMT* with its native promoter and 5′/3′ UTRs fused to *SYFP2* in the *pGEM-T Easy* backbone.

**(b–c)** PCR amplification of the 5′ UTR–*PCKMT* fragment (6,746 bp), SYFP2 (755 bp), and 3′ UTR (516 bp).

**(d)** Fusion PCR joining *SYFP2* with the 3′ UTR (1,235 bp).

**(e–f)** Analytical PCR confirming correct *pGEMT-PCKMT-SYFP2* intermediate (3,119 bp) and full-length construct (7,913 bp).

**(g)** Insertion PCR of PCKMT$^{com}$ -SYFP2 (3212bp) into loxP*pckmt*-Halo/*frm1*-3HA background strain.

**(h–i)** Generation of PCKMT catalytic-mutant constructs(h). Analytical PCR confirming correct mutant fragment sizes of $PCKMT^{H542A}$ (1,093 bp) mutants(i). All verified constructs were sequence-confirmed before transfection.

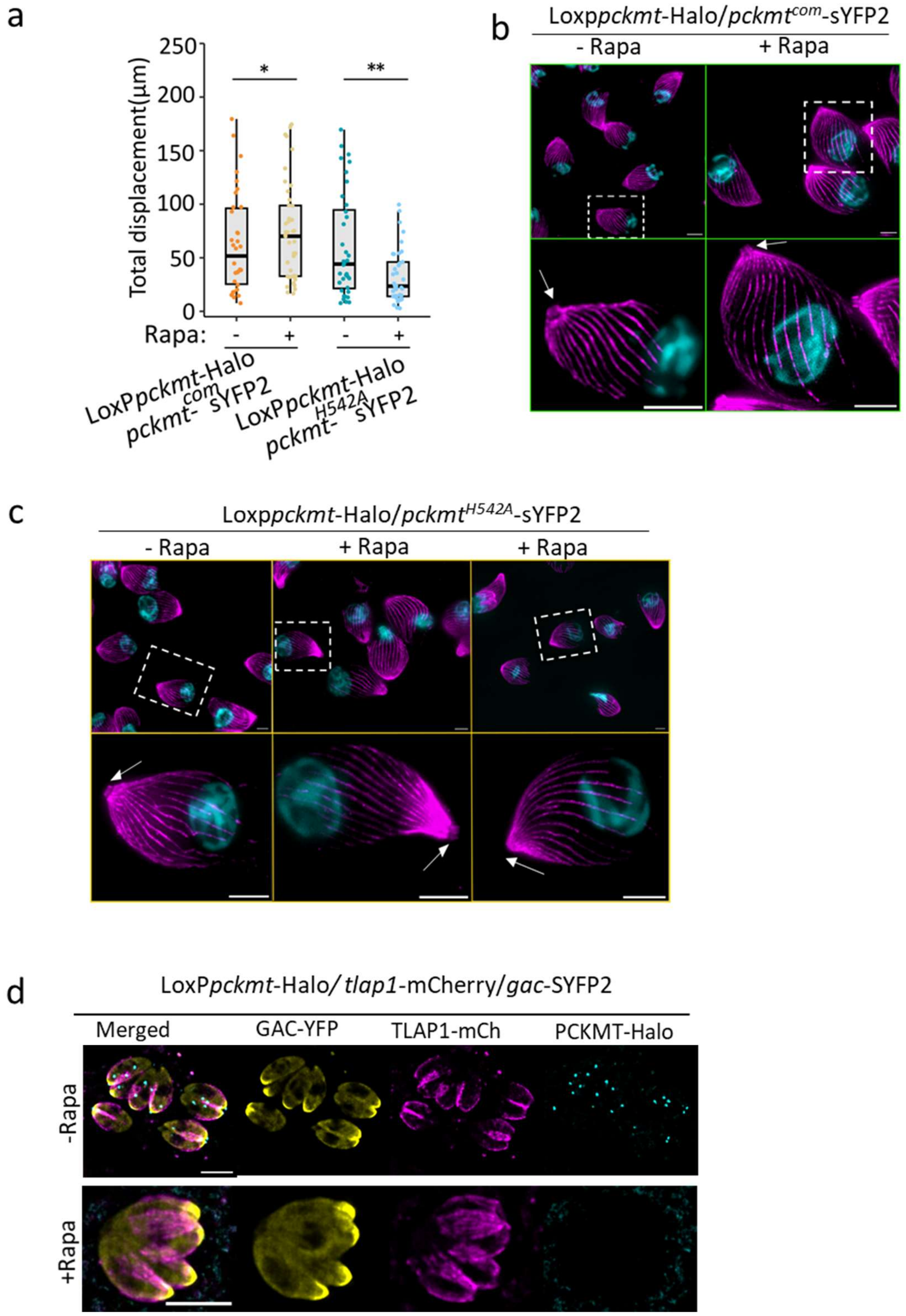


**Supplementary Fig. 6 | Catalytic activity of PCKMT is required for anchoring FRM1 and initiating parasite motility.**

**(a)** Quantification of gliding motility in *T. gondii* tachyzoites expressing wild-type PCKMT or the catalytic mutant $\mathrm{PCKMT}^{H542A}$. Gliding distances were measured from time-lapse recordings using ImageJ. Each point represents an individual parasite (≥9 per replicate, three biological replicates); bars indicate mean ± SD. Statistical analysis was performed using an unpaired, two-tailed Student's *t*-test (*$p < 0.01$, $p < 0.05$).

**(b)** Representative images of conoid extrusion in complemented parasites (PCKMT$^{com}$). Rapamycin-induced excision of endogenous *pckmt* caused a conoid extrusion defect, which was rescued by the expression of PCKMT$^{com}$**.**
**(c)** Representative images of conoid extrusion in the catalytic mutant PCKMT$^{H542A}$. The point mutation failed to restore conoid protrusion following *pckmt* excision. Scale bar, 5 μm.
**(d)** Endogenous tagging of GAC in the conditional loxP*pckmt* strain shows that GAC remains unaffected by PCKMT loss in intracellular parasites, indicating a unidirectional dependency.

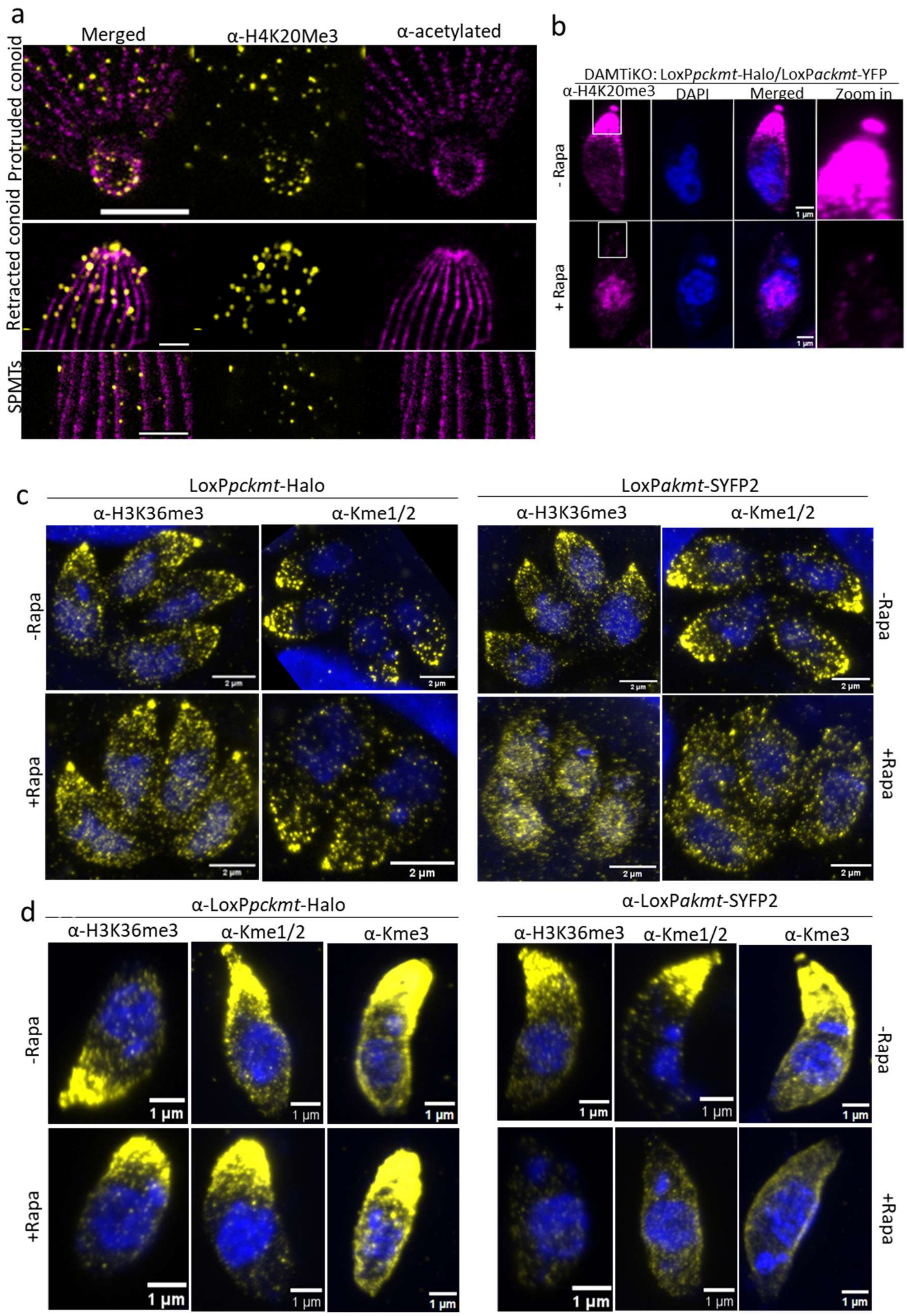


**Supplementary Fig. 7 | Apical methylation resolved by U-ExM/STED reveals AKMT as the dominant lysine methyltransferase, with minimal contribution from PCKMT.**

**(a)** U-ExM combined with STED further enhances the resolution of the methylation signal, preserving distinct labelling at the preconoidal rings and along microtubules after expansion. Scale bar, 2 µm.

**(b)** Immunofluorescence analysis of methylation using anti-H4K20me3 antibody in parasites induced with and without rapamycin (Rapa). Double knockout (DAMTiKO; AKMT/PCKMT). Loss of AKMT abolished the H4K20me3 signal at the apical complex, and the double knockout mirrored this phenotype, confirming AKMT as the dominant methyltransferase responsible for this modification.
**(c)** Intracellular tachyzoites immunolabelled with anti-H3K36me3 and anti-lysine mono/di-methyl (Kme1/2) antibodies. Bulk methylation levels decrease markedly upon AKMT depletion, whereas PCKMT depletion shows no detectable change.
**(d)** Extracellular tachyzoites stained with anti-H3K36me3, anti-Kme1/2, and pan-methyl-lysine antibodies display similar patterns, confirming AKMT as the dominant methyltransferase responsible for global methylation signals**.**

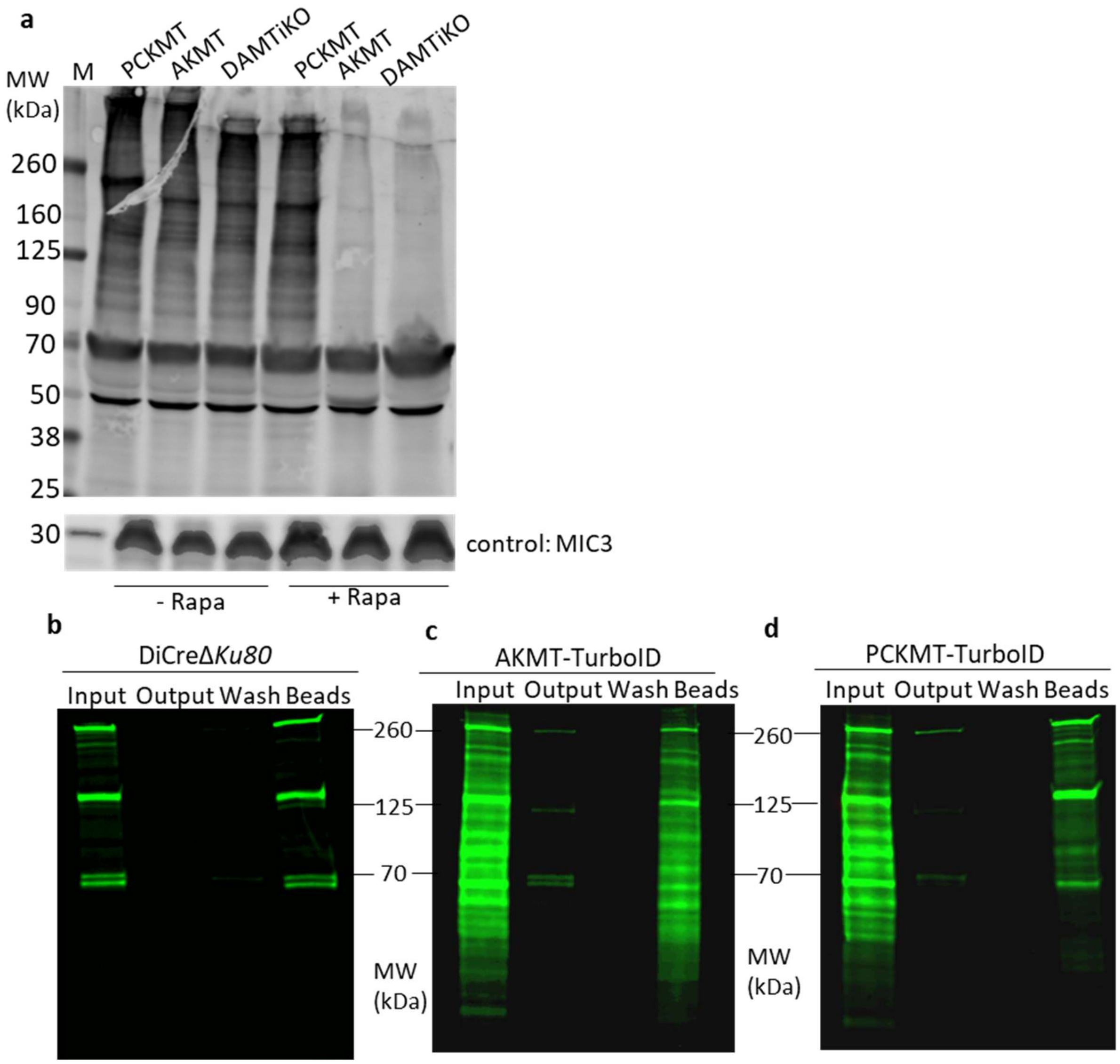


**Supplementary Fig. 8 | Validation of global and proximity-dependent methylation detection in *T. gondii*.**

**(a)** Western blot analysis of total parasite extracts from the indicated strains—*loxPpckmt-Halo*, *loxPakmt-YFP*, and DAMT-iKO—probed with anti-H4K20me3 antibody to assess global lysine methylation. Equivalent loading was confirmed by probing for GAP45.

**(b–d)** Streptavidin-based detection of biotinylated proteins following TurboID proximity labelling. Parasite lysates were resolved by SDS-PAGE and blotted with streptavidin conjugated to infrared dye (IR-800) for detection on a LI-COR Odyssey system.

**(b)** Parental DiCreΔKu80 strain (negative control).

**(c)** AKMT–TurboID parasites showing specific biotinylation of apical and cytosolic proteins.

**(d)** PCKMT–TurboID parasites showing enrichment of preconoidal and apical complex-associated proteins.

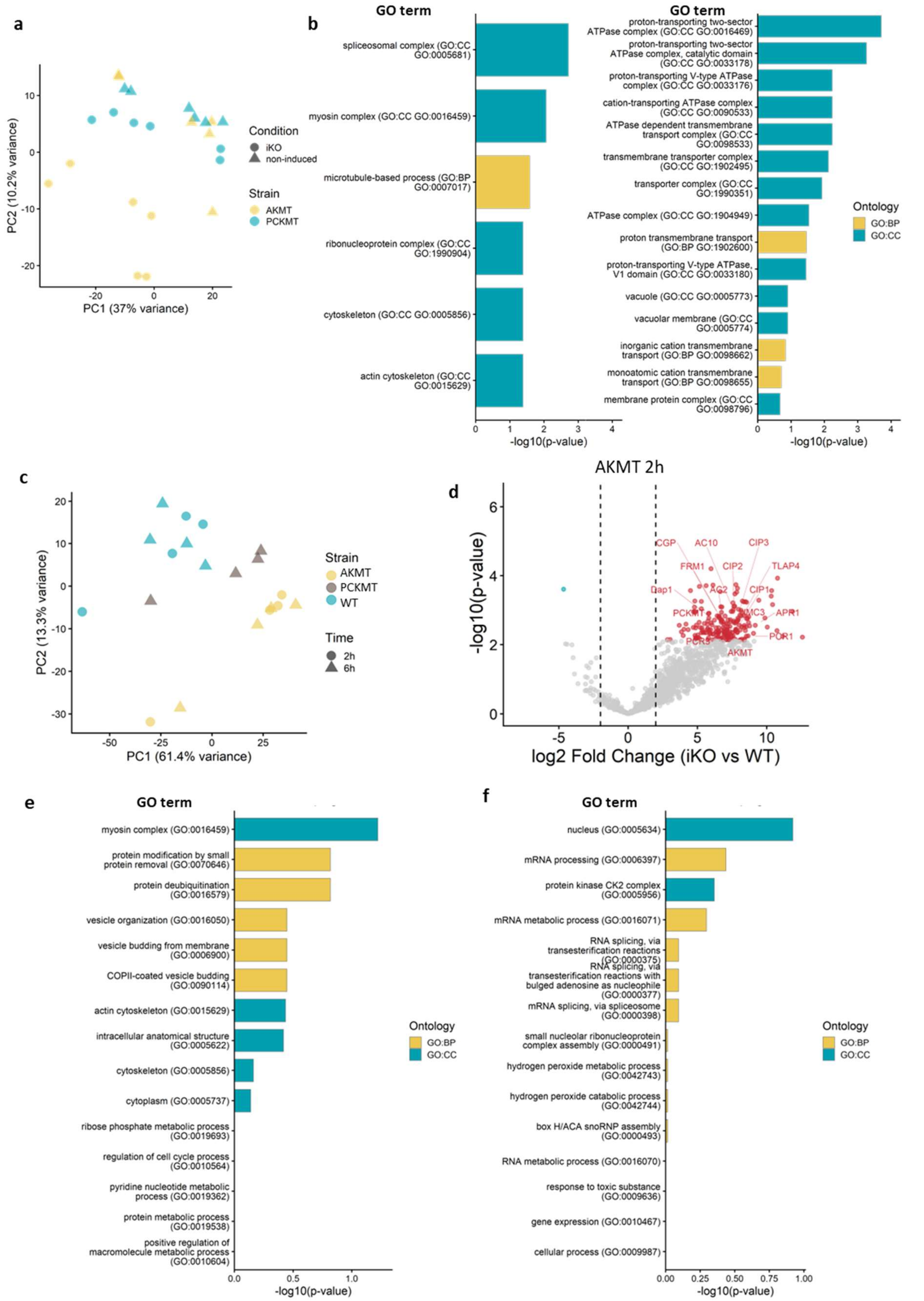


**Supplementary Fig. 9 | Distinct methylation and proximity-labelling signatures define PCKMT and AKMT functional networks in *T. gondii*.**

**(a)** Principal component analysis (PCA) of proteomic features from methylome-enriched samples of *T. gondii* PCKMT- and AKMT-knockout strains. Each point represents one replicate positioned by its scores on PC1 (37% variance) and PC2 (10.2% variance). Colours denote strain (AKMT, yellow; PCKMT, cyan); shapes denote condition (non-induced, circles; iKO, triangles). Tight clustering within groups indicates high reproducibility, while clear separation along PC1/PC2 reflects strain- and induction-specific effects.

**(b)** GO term enrichment analysis of methylation-enriched proteomes. For AKMT (left graph), significantly decreased proteins in the AKMT-iKO versus wild-type were analysed, revealing enrichment of terms associated with cytoskeletal organisation, pellicle structure, and motility-related processes. Due to the limited number of significantly altered proteins in PCKMT-iKO (right graph), GO analysis was performed on proteins with a $\log_2$ fold change $\leq -2$, yielding no meaningful enrichment.

**(c)** Principal component analysis (PCA) of *T. gondii* TurboID proteomes for PCKMT and AKMT. Each point represents one sample positioned by PC1 (37%) and PC2 (10.2%) scores. Colour denotes strain (AKMT, yellow; PCKMT, cyan); shape denotes condition (non-induced, circles; biotin-induced, triangles).

**(d)** Volcano plot of AKMT–TurboID enrichment after 2 h of biotin induction. Cytoskeletal proteins, including FRM1, DAP1, and PCKMT, are among the enriched interactors.

**(e-f)** Gene Ontology (GO) enrichment analysis of TurboID datasets. AKMT interactome shows enrichment for cytoskeletal and vesicle organisation terms **(e)**. PCKMT interactome shows enrichment for nuclear and RNA-processing functions **(f).** GO Biological Process (GO: BP) terms are shown in yellow; GO Cellular Component (GO: CC) in blue.

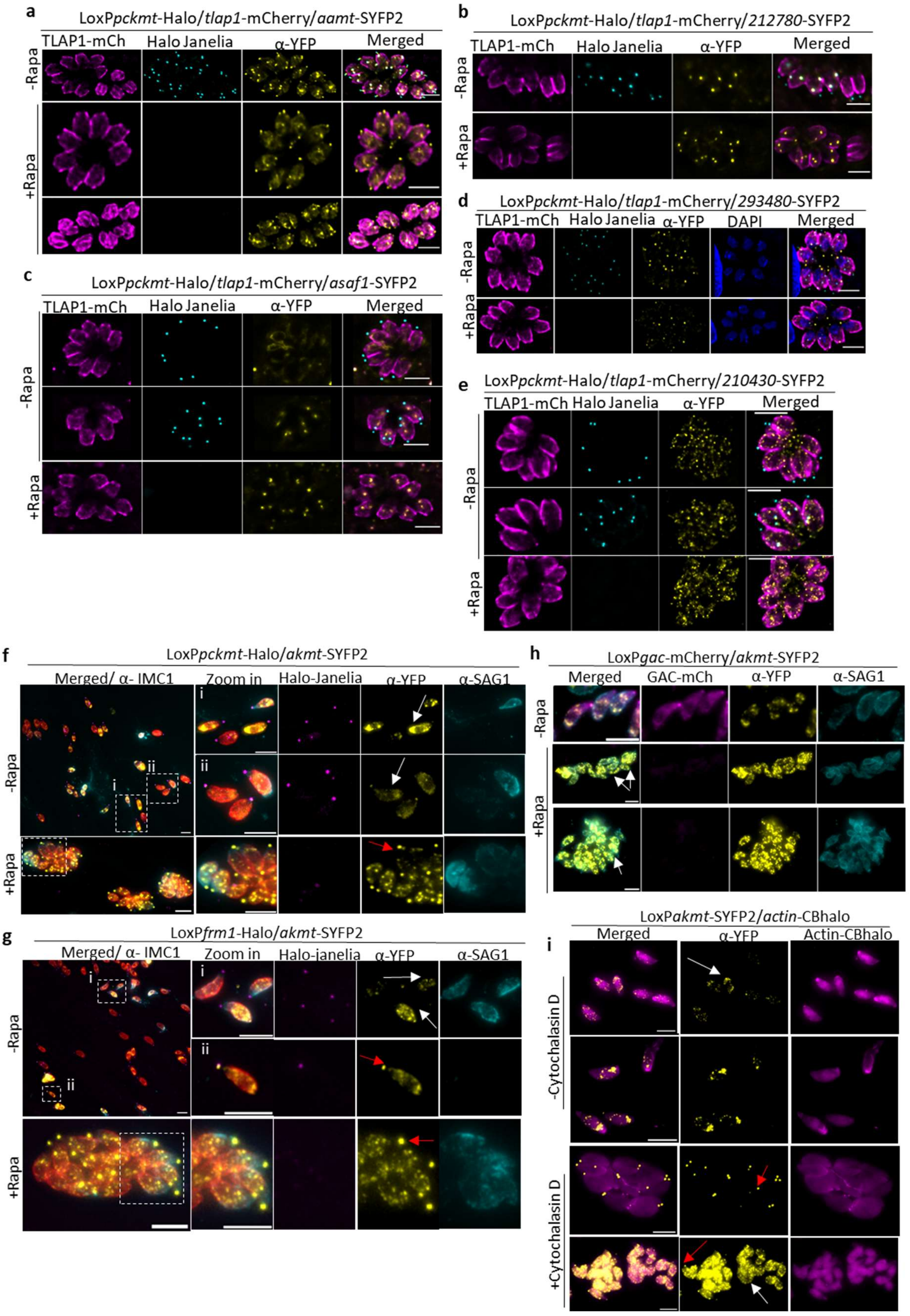


**Supplementary Fig. 10 | PCKMT depletion selectively alters apical protein organisation and modulates AKMT relocalisation dynamics.**

**(a–e)** Endogenous tagging of candidate proteins with YFP in the loxP*pckmt* background. Immunofluorescence assays revealed no detectable changes in localisation following PCKMT depletion.

Scale bar, 5 µm. **(a)** AAMT. **(b)** TGGT1_212780 (hypothetical protein). **(c)** TGGT1_238170 (ASAF1). **(d)** TGGT1_293480 (MoeA N-terminal region-containing protein). **(e)** TGGT1_210430 (DNAJ domain-containing protein).

**(f-i)** Overview images of AKMT translocation shown in Fig. 6a-d. Egress was induced with 4µM CI for 5min. Anti-SAG1 antibody (cyan) was employed to mark the extracellular parasites and vacuoles that had lysed the PVM but had not successfully egressed. IMC(Red) marks all parasites. **(f)** AKMT translocation in loxP*pckmt*-Halo parasites. Parasites were pre-induced ±Rapa for at least 54 h. AKMT re-localises to the cytosol (white arrow). Scale bar: 5 µm. (i) indicates that AKMT sometimes has a strong cytosolic signal in intracellular parasites. (ii) Other times, AKMT has a lower cytosolic signal. **(g)** AKMT translocation in LoxP*frm1*-Halo. (i) Representative image shows the different AKMT intensity in the cytosolic and extracellular parasites (white arrow). (ii) The representative image shows that after egress, the newly invaded parasite has an AKMT signal at the conoid. Scale bar: 5 µm. **(h)** AKMT translocation in loxP*gac*-mCherry (white arrow). Scale bar: 5 µm. **(e)** Parasites were pre-treated with 5µM Cytochalasin D (CytD) for 1h before egress induction. F-actin was labelled with Chromobody-Halo (CbHalo). In some cases, relocalisation of AKMT was observed in the cytosol (white arrow), probably coinciding with a poor abolition of actin dynamics in that vacuole. Red arrow: AKMT is still at the conoid. Scale bar: 5 µm.
